# Markov State Models and Perturbation-Based Approaches Reveal Distinct Dynamic Signatures and Hidden Allosteric Pockets in the Emerging SARS-Cov-2 Spike Omicron Variants Complexes with the Host Receptor: The Interplay of Dynamics and Convergent Evolution Modulates Allostery and Functional Mechanisms

**DOI:** 10.1101/2023.05.20.541592

**Authors:** Sian Xiao, Mohammed Alshahrani, Grace Gupta, Peng Tao, Gennady Verkhivker

## Abstract

The new generation of SARS-CoV-2 Omicron variants displayed a significant growth advantage and the increased viral fitness by acquiring convergent mutations, suggesting that the immune pressure can promote convergent evolution leading to the sudden acceleration of SARS-CoV-2 evolution. In the current study, we combined structural modeling, extensive microsecond MD simulations and Markov state models to characterize conformational landscapes and identify specific dynamic signatures of the SARS-CoV-2 spike complexes with the host receptor ACE2 for the recently emerged highly transmissible XBB.1, XBB.1.5, BQ.1, and BQ.1.1 Omicron variants. Microsecond simulations and Markovian modeling provided a detailed characterization of the conformational landscapes and revealed the increased thermodynamic stabilization of the XBB.1.5 subvariant which is contrasted to more dynamic BQ.1 and BQ.1.1 subvariants. Despite considerable structural similarities, Omicron mutations can induce unique dynamic signatures and specific distributions of conformational states. The results suggested that variant-specific changes of conformational mobility in the functional interfacial loops of the spike receptor binding domain can be fine-tuned through cross-talk between convergent mutations thereby providing an evolutionary path for modulation of immune escape. By combining atomistic simulations and Markovian modeling analysis with perturbation-based approaches, we determined important complementary roles of convergent mutation sites as effectors and receivers of allosteric signaling involved in modulating conformational plasticity at the binding interface and regulating allosteric responses. This study also characterized the dynamics-induced evolution of allosteric pockets in the Omicron complexes that revealed hidden allosteric pockets and suggested that convergent mutation sites could control evolution and distribution of allosteric pockets through modulation of conformational plasticity in the flexible adaptable regions. Through integrative computational approaches, this investigation provides a systematic analysis and comparison of the effects of Omicron subvariants on conformational dynamics and allosteric signaling in the complexes with the ACE2 receptor.

**For Table of Contents Use Only:** 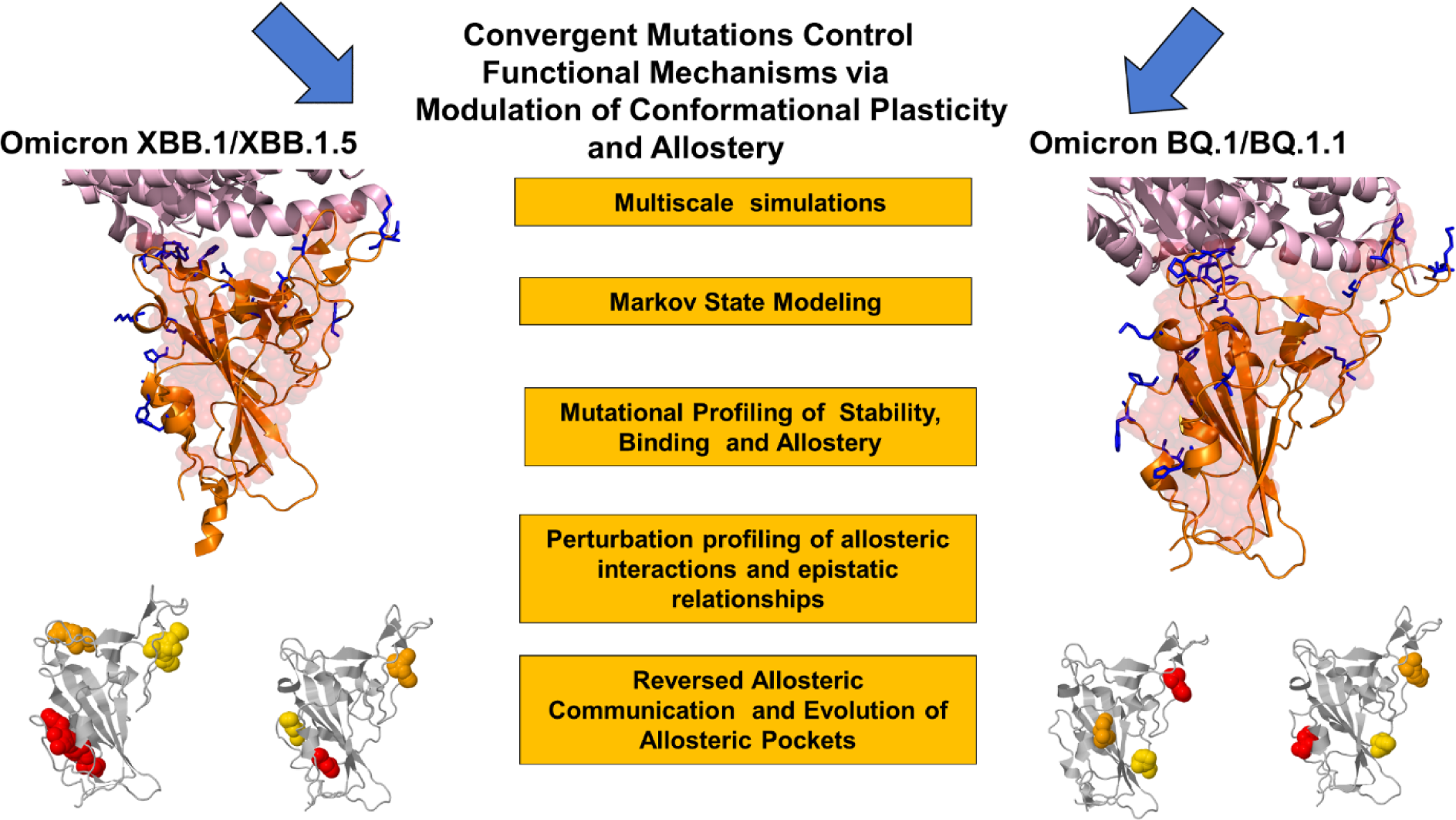

## Introduction

A vast number of structural and biochemical studies have extensively explored the mechanisms underlying SARS-CoV-2 infection.^1-9^ These investigations have unequivocally demonstrated the critical significance of the viral spike (S) glycoprotein in various aspects of the virus’s life cycle, including virus transmission and immune evasion. The S glycoprotein has been identified as a key player in facilitating virus entry into host cells, allowing for viral replication and dissemination. Additionally, it plays a crucial role in evading the host immune response by employing various strategies, such as shielding important epitopes and modulating immune recognition.^1-9^ The architecture of the S protein of SARS-CoV-2 is intricate, comprising of a N-terminal S1 subunit and a C-terminal S2 subunit that is structurally rigid. The S1 subunit undergoes functional movements that involve conformational changes between the closed and open S states. These transitions are facilitated by coordinated global movements of the S1 subunit, which includes the N-terminal domain (NTD), receptor-binding domain (RBD), and two structurally conserved subdomains, SD1 and SD2. Together, these components dictate the S protein’s structural and dynamic response to binding partners and the ACE2 host receptor.^10-15^ Structural and biophysical studies have provided extensive characterization of the thermodynamic and kinetic properties of the functional SARS-CoV-2 S trimer.^16-18^ These studies have focused on exploring the complex interplay between subdomain movements and long-range interactions that link the S1 and S2 subunits to regulate the equilibrium and population-shifts of the RBD between the open (up) and closed (down) conformations. The RBD serves as the primary site of interaction between the S protein and the ACE2 receptor on host cells and the population-shifts of the RBD between open and closed conformations directly impact the accessibility of the ACE2 binding site. The exposure of the RBD in the open conformation can enhance the interaction between the S protein and ACE2, resulting in a higher binding affinity while when the RBD is in the closed conformation, the binding affinity may be weakened or inhibited. This regulatory mechanism allows the S protein to finely modulate its interaction with ACE2 and determine the efficiency of viral entry into host cells.^16-18^

Cryo-electron microscopy (cryo-EM) and X-ray structural studies have been instrumental in elucidating structural characteristics of the SARS-CoV-2 S protein variants of concern (VOCs) in different functional states, as well as their complexes with antibodies.^19-28^ These investigations have unveiled a remarkable diversity of binding epitopes and highlighted the versatility of the S protein’s binding mechanisms when interacting with various classes of antibodies. Biophysical studies using the differential scanning fluorimetry (DSF) assay have investigated the protein stability of SARS-CoV-2 Wu-Hu-1, Delta, and Omicron variants showing that the S Omicron BA.1 variant exhibits the lower inflection temperature and a reduced protein stability for the S Omicron BA.1 variant as compared to the S Wu-Hu-1 and S Delta variants, while the BA.2 RBD was found to be more stable than BA.1 variant.^29-32^ In contrast to the conformational changes observed in the original Wu-Hu-1 strain and other VOCs upon ACE2 binding, the Omicron S protein exhibits a distinct behavior showing a preference for the “one-RBD-up” conformation both before and after ACE2 binding.^33-35^ The Omicron BA.2 subvariants of SARS-CoV-2 have been linked to the increased transmissibility, and especially significant antibody and vaccine evasion capabilities.^36-40^ Studies examining the binding affinities of Omicron BA.1.1 and BA.2 with the ACE2 receptor revealed that these variants exhibit stronger binding affinity compared to the BA.3 and BA.1 variants, suggesting that these variants may have a higher potential for viral entry into host cells.^41-43^ At the same time, surface plasmon resonance (SPR) studies showed that the Omicron BA.4/5 RBD have only marginally greater binding affinity for ACE2 as compared to the ancestral Wu-Hu-1 strain and the BA.1 variants.^44^ The cryo-EM structures and biochemical analysis of the S trimers for BA.1, BA.2, BA.3, and BA.4/BA.5 subvariants confirmed that the increased transmissibility of the BA.2 variant can be driven by a combination of the enhanced ACE2 binding affinity and stronger antibody evasion ad compared to the other Omicron sublineages.^45,46^ The cryo-EM conformations of the BA.2.75 S trimer in the open and closed forms, as well as the structures of the open BA.2.75 complexes with ACE2, have shown that the BA.2.75 is the most stable among the Omicron variants, exhibiting ∼ 4-6-fold increased binding affinity as compared to other Omicron variants.^47,48^

The BA.2 variant generated a wave that led to “second-generation” newly emerging variants display significant growth advantage including BA.2.3.20, BA.2.75, BQ.1, BQ.1.1 and especially XBB.1, XBB.1.5, and XBB.1.6 subvariants that demonstrated the increased transmissibility and the ability to outcompete other strains in the rate infectivity and spread.^49,50^ BQ.1 is a direct descendant of BA.5, with additional spike mutations in some key antigenic sites (K444T and N460K). Its first descendant, BQ.1.1 carries a further additional mutation R346T. The XBB.1 subvariant is derived from the BA.2 lineage and emerged by recombination of two co-circulating BA.2 lineages (BJ.1 and BM.1.1.1). The XBB.1.5 subvariant harbors F486P modification involving a two nucleotide substitution and, while it is equally immune evasive to closely related XBB.1 with F486S mutation, XBB.1.5 exhibits growth advantage due to the higher ACE2 binding affinity from a single S486P mutation that can restore most of the favorable hydrophobic contacts.^51-53^ These studies suggested that R346T and L368I substitutions may enhance binding affinity, while F486S in XBB.1 leads to the decreased ACE2 binding affinity. Biochemical investigations also demonstrated that suggested that the binding affinity of XBB RBD is enhanced compared to BA.2 but is weaker than that of BA.2.75 variant, while the ACE2 binding affinities of BQ.1 and BQ.1.1 were comparable to that of BA.4/5 variants.^53,54^ The binding affinity of the XBB.1.5 RBD to ACE2 was only slightly weaker to that of BA.2.75 but significantly stronger than that of XBB.1 and BQ.1.1 variants.^55^ The neutralizing activity against XBB.1.5 was considerably lower than that against the ancestral strain and BA.2, while similar immune evasion potential was observed for XBB.1 and XBB.1.5 variants.^56,57^ These studies concluded that a high transmissibility and rapid surge of the newly emerged variants may be primarily due to a significant immune evasion potential and the enhanced fitness of the highly favorable ACE2 binding affinity, which is comparable only to the BA.2.75 variant.

Since March 2023, sublineages of the XBB variant harboring the F486P substitution (XBB.1.5 and XBB.1.9) dominated worldwide (https://nextstrain.org/ncov/gisaid/global/6m). Very recently, an XBB sublineage XBB.1.16 RBD emerged and was detected in various countries including the US. Compared to XBB.1.5, XBB.1.16 has E180V in the NTD and T478R in the XBB.1.16 has a greater growth advantage in the human population compared to XBB.1 and XBB.1.5, while the immune evasion potential of XBB.1.16 is comparable to XBB.1 and XBB.1.5 subvariants.^58^ The newly emerging variants display substantial growth advantages over previous Omicron variants, and some RBD residues (R346, K356, K444, V445, G446, N450, L452, N460, F486, F490, R493 and S494) are mutated in at least five independent Omicron sublineages that exhibited a high growth advantage.^59^ BQ.1.1 subvariant bears all five recent convergent mutations: R346T, K444T, L452R, N460K, and F486V. Evolutionary studies suggested that sublineages descending from BA.5, including BQ.1 and BQ.1.1 convergently increased viral fitness by consecutively acquiring substitutions at the R346, N460, and K444 residues and it was proposed that BQ.1.1 increased the binding affinity to human ACE2 during evolution from BA.5 variant.^60^ Consistent with previous studies from the same group^51,52^ L452R and N460K substitution can increase the ACE2 binding affinity while K444T substitution significantly decreased ACE2 binding affinity.^60^ It was suggested that acquiring L452R and N460K substitutions that potentially increase ACE2 binding ability can allow for the increased substitution frequency at the convergent sites to modulate immune escape capacity. Indeed, latest studies showed that several convergent mutations present in the BA.2 or BA.5 subvariants including R346T mutation found in BA.2.75.2, along with K444T and N460K substitutions are associated with escape from monoclonal antibodies and from vaccine-induced antibodies.^61,62^ The patterns of convergent evolution in the SARS-CoV-2 variants was reviewed recently using analysis of the mutations which emerged at least three times independently in different linages.^63^ The convergently evolved mutations tend to cluster in specific locations of NTD and RBD, and spatial clustering under convergent evolution can create synergic epistatic effects between RBD mutations.^64,65^ Additionally, convergent mutations N439K, N440K, L452R, T478R/K and E484K may have evolved to carry positive charges and enhance electrostatic complementarity with ACE2. Importantly, it was concluded that balancing ACE2 binding and immune escape fitness tradeoffs can shape up the RBD evolution and patterns of convergent mutations. By examining possible mechanisms driving the emergence of convergent RBD mutations it was suggested that the immune pressure on the RBD becomes increasingly focused and promotes convergent evolution, explaining the observed sudden acceleration of SARS-CoV-2 RBD evolution and the convergence pattern.^66^

Computer simulations of the SARS-CoV-2 S proteins have allowed for a better understanding of the molecular mechanisms of SARS-CoV-2 S protein and receptor binding. Molecular dynamics (MD) simulations of the full-length SARS-CoV-2 S glycoprotein embedded in the viral membrane, with a complete glycosylation profile provided detailed characterization of the conformational landscapes of the S proteins in the physiological environment.^67-70^ Using distributed cloud-based computing, large scale MD simulations of the viral proteome observed dramatic opening of the S protein complex, predicting the existence of several cryptic epitopes in the S protein.^71^ MD simulations of the S protein in solution and targeted simulations of conformational changes between the open and closed S forms revealed the key electrostatic interdomain interactions mediating the protein stability and kinetics of the functional spike states.^72^ Using the replica-exchange MD simulations, conformational landscapes of the full-length S protein trimers were investigated, unveiling hidden functional intermediates along with open-closed transition pathways and previously unknown cryptic pockets.^73^ Using MD simulations with a full representation of glycan moieties, conformational changes in a complete membrane-embedded SARS-CoV-2 spike glycoprotein were probed to accurately describe pocket dynamics leading to discovery of cryptic pockets.^74^ The linkage between conformational dynamics and allosteric modulation of SARS-CoV-2 S in the absence or presence of ligands was established in an smFRET imaging assay, showing presence of long-range allosteric control of the RBD equilibrium, which in turn regulates the exposure of the binding site and antibody binding.^75^ Our studies used integrative computational modeling approaches to reveal that the S protein could function as an allosteric regulatory machinery controlled by stable allosteric hotspots acting as drives drivers and regulators of spike activity.^76-82^

A comparative analysis of the conformational dynamics, electrostatics, protein stability and allostery in the different functional states of S trimers for BA.1, BA.2, and BA.2.75 variants showed that Omicron mutations may evolve by targeting vulnerable sites of conformational adaptability to elicit immune escape, while maintaining their control on balancing protein stability and functional fitness through robust allosteric communications with the stability hotspots.^83^ By combining atomistic simulations and a community-based network model of epistatic couplings we found that convergent Omicron mutations such as G446S (BA.2.75, BA.2.75.2, XBB), F486V (BA.4, BA.5, BQ.1, BQ.1.1), F486S, F490S (XBB.1), F486P (XBB.1.5) can display epistatic relationships with the major stability and binding affinity hotspots which may allow for the observed broad antibody resistance.^84^

In the current study, we combine multiple microsecond MD simulations and MSM analysis to characterize conformational landscapes and identify specific dynamic signatures of the SARS-CoV-2 S RBD-ACE2 complexes for the recently emerged XBB.1, XBB.1.5, BQ.1, and BQ.1.1 Omicron variants. Microsecond atomistic simulations and MSM analysis unveil unique dynamic signatures and provide a detailed characterization of the conformational landscapes showing the increased thermodynamic stabilization of the XBB.1.5 subvariant which is contrasted to more dynamic BQ.1 and BQ.1.1 subvariants. Using a comparative MSM analysis we discover variant-specific changes of conformational mobility in the flexible RBD loops 470-491 and 440-452 containing critical convergent mutation sites that may allow for modulation of conformational plasticity and immune escape. By combining MD simulations and MSM analysis with perturbation-based response scanning approaches, we also examine mechanisms of long-range dynamic couplings and allosteric communications in the Omicron RBD-ACE2 complexes. Based on premise that allostery is linked with the evolvability of proteins, we explore the relationship between allosteric interactions and patterns of convergent Omicron mutations, showing how convergent mutations can potentiate conformational plasticity and modulate allosteric responses in the RBD to binding and immune escape. Through perturbation response scanning analysis, our study reveals important complementary roles of convergent mutation sites as effectors/regulators and sensors/receivers of S-RBD plasticity allosteric signal transmission in the RBD-ACE2 complexes. We also explore our recently developed method for fast and accurate allosteric site prediction PASSer (Protein Allosteric Sites Server) (https://passer.smu.edu)^85-88^ to characterize the effect of dynamics on the distribution of allosteric pockets in the Omicron subvariants. The results show that variant-specific redistribution of macrostates preserves the experimentally known allosteric pocket on the RBD while allowing for the emergence of hidden allosteric pockets that are anchored by convergent mutation sites K444T and L452R. Through integrative computational approaches, this study provides a systematic analysis and comparison of the effects of Omicron subvariants on conformational dynamics and allosteric signaling in the complexes with the ACE2 receptor.

## Materials and Methods

### Structural modeling and refinement

The crystal structures of the BA.2 RBD-ACE2 (pdb id 7XB0), and BA.5 RBD-ACE2 complexes (pdb id 7XWA) were obtained from the Protein Data Bank.^89^ During structure preparation stage, protein residues in the crystal structures were inspected for missing residues and protons. Hydrogen atoms and missing residues were initially added and assigned according to the WHATIF program web interface.^90^ The missing loops in the studied cryo-EM structures of the SARS-CoV-2 S protein were reconstructed and optimized using template-based loop prediction approach ArchPRED.^91^ The side chain rotamers were refined and optimized by SCWRL4 tool.^92^ The protein structures were then optimized using atomic-level energy minimization with composite physics and knowledge-based force fields implemented in the 3Drefine method.^93,94^ The refined structural models of the XBB.1 RBD-ACE2 and XBB.1.5 RBD-ACE2 complexes were obtained based on its paternal BA.2 RBD-ACE2 complex and the models for BQ.1 and BQ.1.1 complexes were constructed based on the paternal BA.5 RBD-ACE2 complex. MutaBind2 approach was employed to introduce mutations and optimize structural models for XBB.1, XBB.1.5, BQ.1 and BQ.1.1 RBD-ACE2 complexes. MutanBind2 utilizes molecular mechanics force fields and fast side-chain optimization algorithms via random forest method.^95,96^ MutaBind2 utilizes FoldX approach^97,98^ to introduce single or multiple point mutations on the crystal structure followed by robust side-chain optimization and multiple rounds of energy minimization using NAMD 2.9 program^99^ with CHARMM36 force field.^100^

### All-Atom Molecular Dynamics Simulations

The CHARMM36 force field^100^ with the TIP3P water model^101^ were employed to perform all-atom MD simulations for each of the Omicron RBD-ACE2 complexes. The structures of the SARS-CoV-2 S-RBD complexes were prepared in Visual Molecular Dynamics (VMD 1.9.3).^102^ The protonation states of titratable residues were determined under neutral pH. The protein systems were solvated in 130 Å × 85 Å × 75 Å water boxes. In each system, sodium and chloride ions were added to maintain an ionic strength of 0.1 M. After energy minimization, the systems were first heated up from 100 to 300 K with a temperature increment of 20 K per 50 picoseconds (ps). Consequently, the systems were subjected to 1.5 nanoseconds (ns) isothermal−isobaric (NPT) equilibrations at 300 K (equilibrium run), followed by 1 microsecond (µs) canonical (NVT) simulations (production run) at 300 K. Snapshots of the production run were saved every 100 ps. In all simulations, the SHAKE constraint was used to constrain bonds associated with hydrogen atoms in the solvent molecules and the proteins.^103^ The nonbonding interactions within 10 Å were calculated explicitly. The Lennard-Jones interactions were smoothed out to zero at 12 Å. The long-range electrostatic interactions were calculated using the particle mesh Ewald method^104^ with a cut-off of 1.0 nm and a fourth order (cubic) interpolation. The simulations were conducted using OpenMM (version 7.6.0).^105^ For each system, MD simulations were conducted three times in parallel to obtain comprehensive sampling. Each individual simulation has 10,000 frames.

### Time-Structure based Independent Component Analysis (tICA)

The tICA method identifis the slowest degrees of freedom and therefore in preserving the kinetic information present in the MD trajectories by maximizing the auto-correlation function.^106-109^ Usng MD trajectories, ***x***(*t*) = (*x*_1_(*t*),…,*x_n_*(*t*)), tICA reduces the dimensionality of the trajectories and to identify hidden key structural changes by decomposing the generalized eigenvalue problem *CF = CFK* where *K* = *diag*(*k*_1_,…,*k_n_*) and *F* = (*f*_1_,…,*f_n_*) are the eigenvalue and eigenvector matrices, respectively; *C* and *C* are the covariance matrix and the time-lagged covariance matrix of the coordinate vector, defined as follows

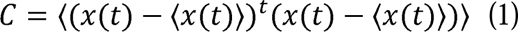

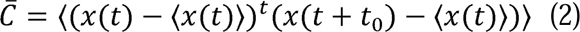

where 〈…〉 denotes the average. In order to obtain a symmetric time-lagged covariance matrix 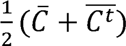 is calculated. The latter step assumes the time reversibility of the process, which is satisfied in MD simulations. The projected vectors of the MD are expressed as:

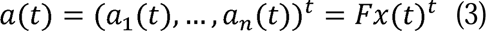

The featurization and dimensionality reduction were performed using the MSMBuilder package.^110^

### Markov State Model

Stochastic Markov state models (MSMs)^111-113^ have become increasingly useful states-and-rates models with the mature and robust software infrastructure^114,115^ for describing the transitions between functional protein states and modeling of allosteric events. In MSM, protein dynamics is modeled as a kinetic process consisting of a series of Markovian transitions between different conformational states at discrete time intervals. A specific time interval, referred to as lag time, needs to be determined to construct transition matrix. First, k-means clustering method is conducted on projected low-dimensional space and each simulation frame is assigned to a macrostate. The transition counting is constructed based on a specific time interval lag time. Macrostates were kinetically clustered based on the Perron-cluster cluster analysis (PCCA++)^116^ and are considered as kinetically separate equilibrium states. The transition matrix and transition probability were calculated to quantify the transition dynamics among macrostates. The corresponding transition probability from state to state is calculated as:

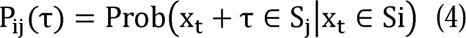

A proper lag time is required for MSM to be Markovian. The value of the lag time and the number of macrostates are selected based on the result of estimated relaxation timescale.^117^ The implied time scale can be calculated using the eigenvalues *λ_i_* in the transition matrix as

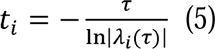

The number of protein metastable states associated with these slow relaxation timescales can be inferred based on the convergence of implied relaxation time scale. These metastable states effectively discretize the conformational landscape. The Markov state model building was conducted using PyEMMA package (v2.5.12).^118^ Based on the transition matrix we obtain implied timescales for transitioning between various regions of phase space and use this information determines the number of metastable states. The number of metastable states also defines the resolution of the model by determining how large a barrier must be in order to divide phase space into multiple states.

### Perturbation Response Scanning

Perturbation Response Scanning (PRS) approach^119-121^ follows the protocol originally proposed by Bahar and colleagues^122,123^ and was described in detail in our previous studies.^124,125^ In this approach, it 3N × 3*N* Hessian matrix ***H*** whose elements represent second derivatives of the potential at the local minimum connect the perturbation forces to the residue displacements. The 3*N*-dimensional vector **Δ*R*** of node displacements in response to 3*N*-dimensional perturbation force follows Hooke’s law ***F = H * ΔR***. A perturbation force is applied to one residue at a time, and the response of the protein system is measured by the displacement vector Δ***R***(*i*) = ***H*^−1^ *F^(i)^*** that is then translated into *N*×*N* PRS matrix. The second derivatives matrix ***H*** is obtained from simulation trajectories for each protein structure, with residues represented by *C_α_* atoms and the deviation of each residue from an average structure was calculated by Δ**R**_*j*_(*t*) = **R**_*j*_(*t*) − 〈**R**_*j*_(*t*)〉, and corresponding covariance matrix C was then calculated by Δ**R**Δ^*T*^. We sequentially perturbed each residue in the SARS-CoV-2 spike structures by applying a total of 250 random forces to each residue to mimic a sphere of randomly selected directions. The displacement changes, Δ***R^i^*** is a *3N-*dimensional vector describing the linear response of the protein and deformation of all the residues. Using the residue displacements upon multiple external force perturbations, we compute the magnitude of the response of residue *k* as 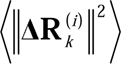 averaged over multiple perturbation forces **F**^(*i*)^, yielding the *ik*^th^ element of the *N*×*N* PRS matrix. The average effect of the perturbed effector site *i* on all other residues is computed by averaging over all sensors (receivers) residues and can be expressed as 〈(Δ***R^i^***)^2^〉_*effector*_. The effector profile determines the global influence of a given residue node on the perturbations in other protein residues and can be used as proxy for detecting allosteric regulatory hotspots in the interaction networks. In turn, the *j* ^th^ column of the PRS matrix describes the sensitivity profile of sensor residue *j* in response to perturbations of all residues and its average is denoted as 〈(Δ**R**^*i*^)^2^〉_*sensor*_. The sensor profile measures the ability of residue *j* to serve as a receiver of dynamic changes in the system.

### Machine Learning Based Prediction of Allosteric Pockets

We employ a machine learning model that is capable of ranking pockets in order of their likelihood to be allosteric sites. FPocket was applied on each protein to detect protein pockets.^126^ For each detected pocket, 19 physical and chemical features are calculated, ranging from pocket volume, solvent accessible surface area to hydrophobicity. We used the LTR algorithm using GBDT and the LambdaMART method. GBDT is a machine learning approach that iteratively learns from decision trees and ensembles of their predictions. Here, we use LightGBM^127^ one of the two popular implementations of GBDT, over XGBoost.^128^ LambdaMART is an LTR method that trains GBDT with the lambdarank loss function. The lambdarank loss function optimizes the value of the normalized discounted cumulative gain (NDCG) for the top *K* cases, and is calculated using discounted cumulative gain (DCG) and ideal discounted cumulative gain (IDCG) as:

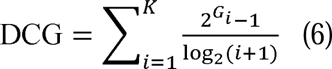

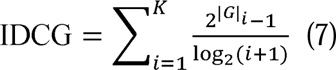

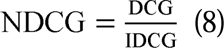

where *G_i_* is the gain (graded relevance value) at position *i* and |*G*|is the ideal ranking.

The LGBMRanker module in the LightGBM package (v3.3.4) was used to implement the LambdaMART algorithm with GBDT as boosting type and lambdarank as the objective function. In the learning-to-rank model, the ASD proteins were randomly split into a training set (80%) and a test set (20%). Five-fold cross validation was performed on the training set for parameter tuning, and the best-performed parameter setting was selected and used on the test set.

## Results and Discussion

### Microsecond MD Simulations of the Omicron RBD-ACE2 Complexes Reveal Distinct Dynamic Signatures of RBD Proteins

We first performed a comparative structural analysis of the Omicron RBD-ACE2 complexes for XBB.1 and XBB.1.5 complexes with the ACE2 receptor (Figure 1). Mutations G339D, S373P, S375F, K417N, N440K, S477N, T478K, E484A, Q493R, Q498R, N501Y, and Y505H in BA.2 are shared with the BA.1 variant, but BA.2 additionally carries S371F, T376A, D405N, and R408S mutations (Table 1). BA.2.75 subvariant has nine additional mutations as compared to BA.2 in the NTD (K147E, W152R, F157L, I210V, and G257S) and RBD (D339H, G446S, N460K, and R493Q) (Table 1). G446S mutation is shared with Omicron BA.1, and R493Q reversed mutation is present in BA.4/BA.5 as well as in XBB.1 and XBB.1.5 subvariants. XBB.1 subvariant is a descendant of BA.2 and recombinant of BA.2.10.1 and BA.2.75 sublineages, featuring NTD mutations V83A, H146Q, Q183E, V213E, G252V and specific RBD mutations (G339H, R346T, L368I, V445P, G446S, N460K, F486S, F490S and reversed R493Q) (Table 1). Many of these RBD mutations are known for their immune evasion functions, including R346T, G446S and F486S.^57^ XBB.1.5 is remarkably similar to XBB.1 with a critical single RBD modification F486P mutation (Table 1). Among the most recent descendant lineages is the Omicron subvariant BQ.1 which is a direct descendant of BA.5, with additional spike mutations in some key antigenic sites (K444T and N460K). Its first descendant, BQ.1.1 carries a further additional mutation (R346T)^130^. BQ.1 has been detected in 65 countries, with an overall prevalence of 6%.

**Figure 1.**
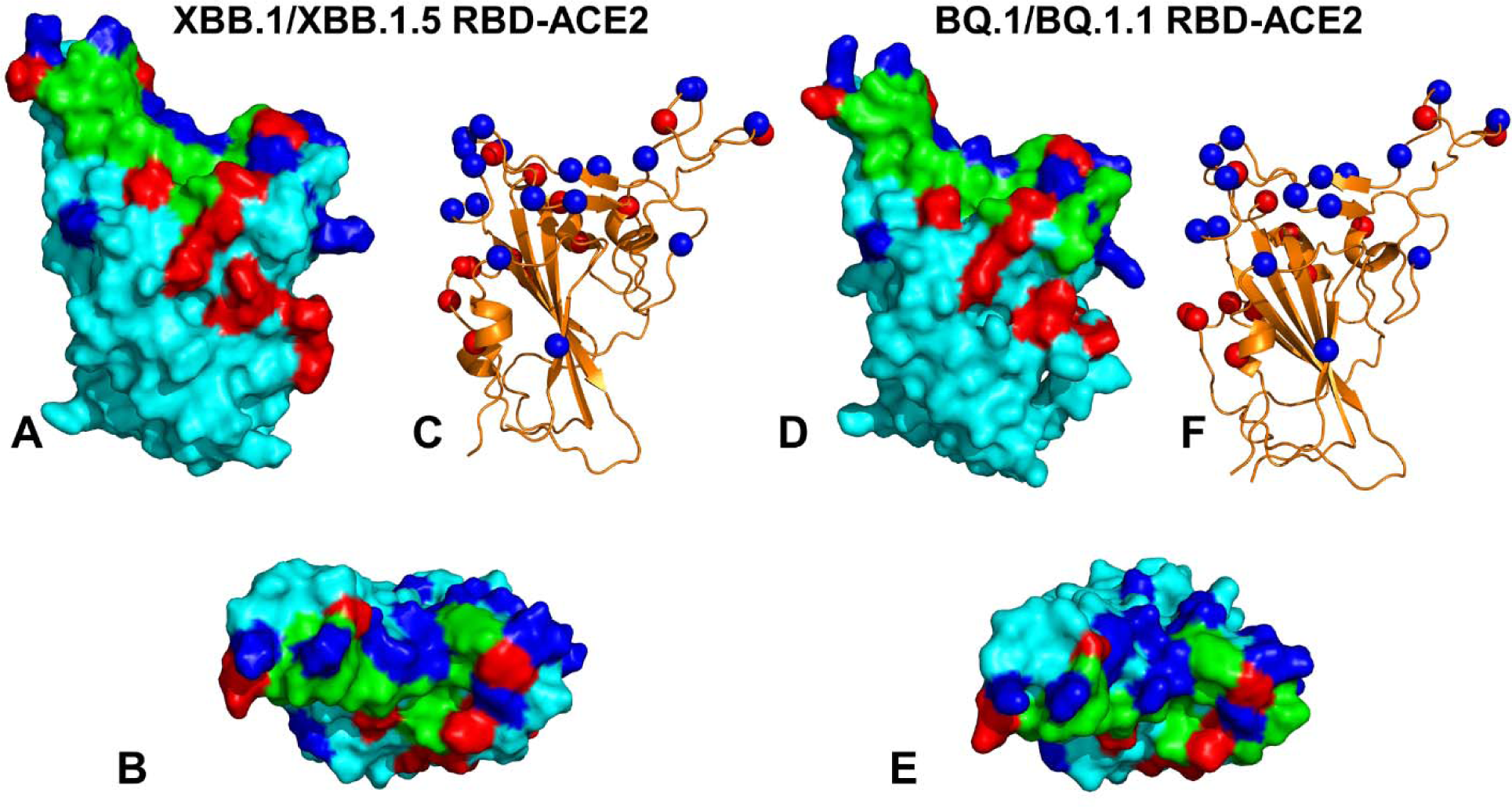
Structural organization and binding epitopes of the SARS-CoV-2-RBD Omicron XBB.1/XBB.1.5 and BQ.1/BQ.1.1 complexes with human ACE enzyme. (A) The optimized modeled structure of the Omicron RBD XBB.1/XBB.1.5-ACE2 complex (only RBD is shown). The RBD is in cyan surface, The binding epitope is in green surface, XBB.1/XBB.1.5 Omicron mutations are in red, and the convergent mutational sites are in blue surface. (B) The top view of the RBD XBB.1/XBB.1.5 from complexes with ACE2. (C) The Omicron RBD XBB.1/XBB.1.5 in ribbon representation. The sites of convergent mutations are in blue spheres. (D) The optimized modeled structure of the Omicron RBD BQ.1/BQ.1.1-ACE2 complex (only RBD is shown). The RBD is in cyan surface, The binding epitope is in green surface, BQ.1/BQ.1.1 Omicron mutations are in red, and the convergent mutational sites (R346T, K356, N440K, K444T, V445, G446, N450, L452R, N460K, T478K, E484A, F486V, F490, Q493, S494, N501Y) are in blue surface. (E) The top view of the RBD BQ.1/BQ.1.1 from complexes with ACE2. (F) The Omicron RBD BQ.1/BQ.1.1 in ribbon representation. The sites of convergent mutations are shown in blue spheres.

**Table 1.**
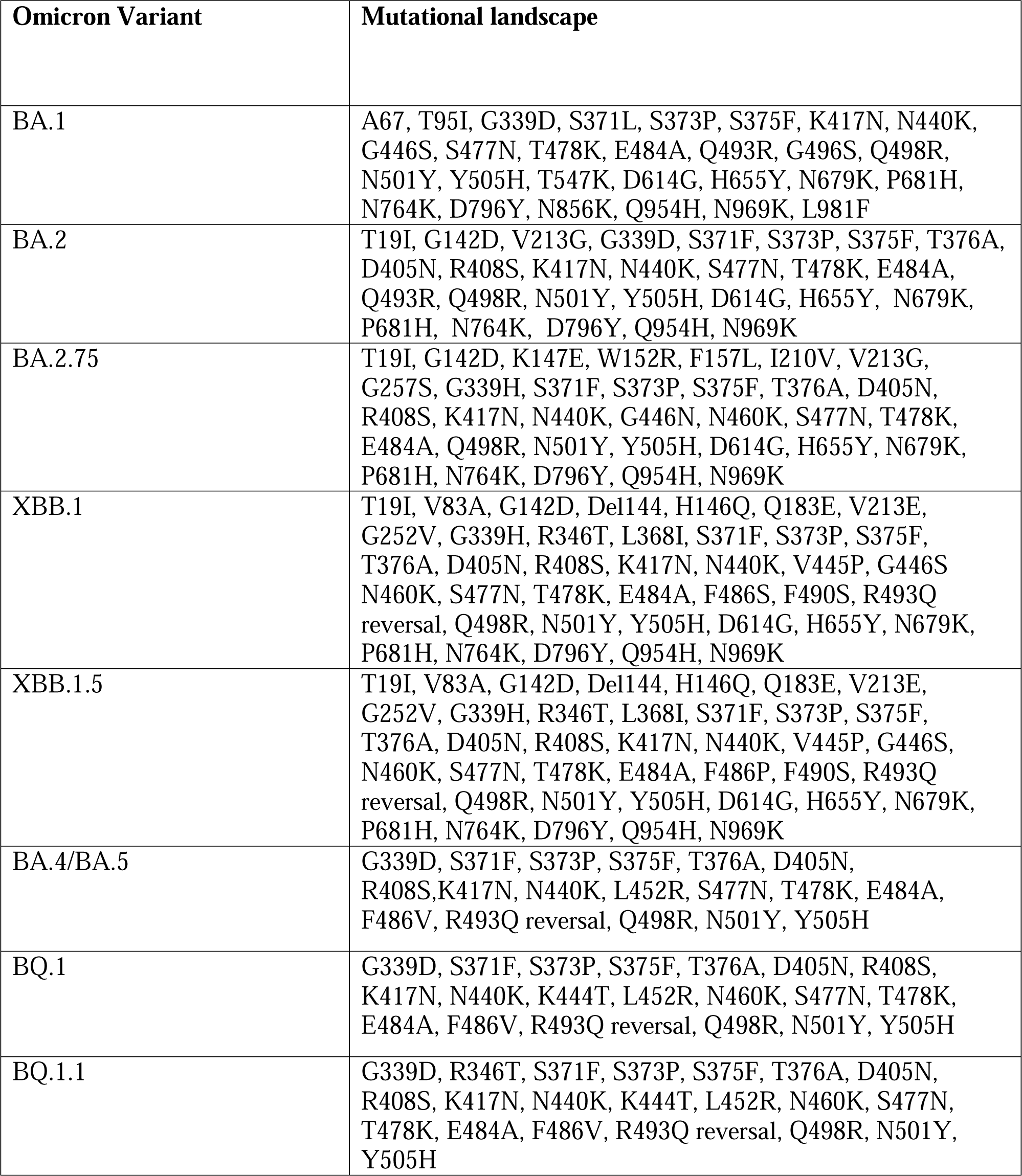
Mutational landscape of the Omicron subvariants in the S-RBD.

Despite considerable mutational differences between newly emerged Omicron variants, structural analysis of the RBD complexes with ACE2 for these variants revealed highly similar RBD conformations and the same binding mode of interactions, rendering overall very minor differences in the crystallographic conformations (Figure 1). We conjectured that the regulatory mechanisms of Omicron variants and functional differences in stability, binding and immune escape can be examined through the lens of the conformational dynamics, reconstruction of the conformational landscape and analysis of the distribution of multiple substates available to these systems. To characterize conformational landscapes and dynamic signatures of the Omicron variants, we conducted multiple independent microsecond MD simulations for each of the studied RBD-ACE2 complexes (Tables 1,2, Figure 2) MD simulations revealed important commonalities and striking differences in the intrinsic conformational dynamics of the RBDs among Omicron variants.

**Figure 2.**
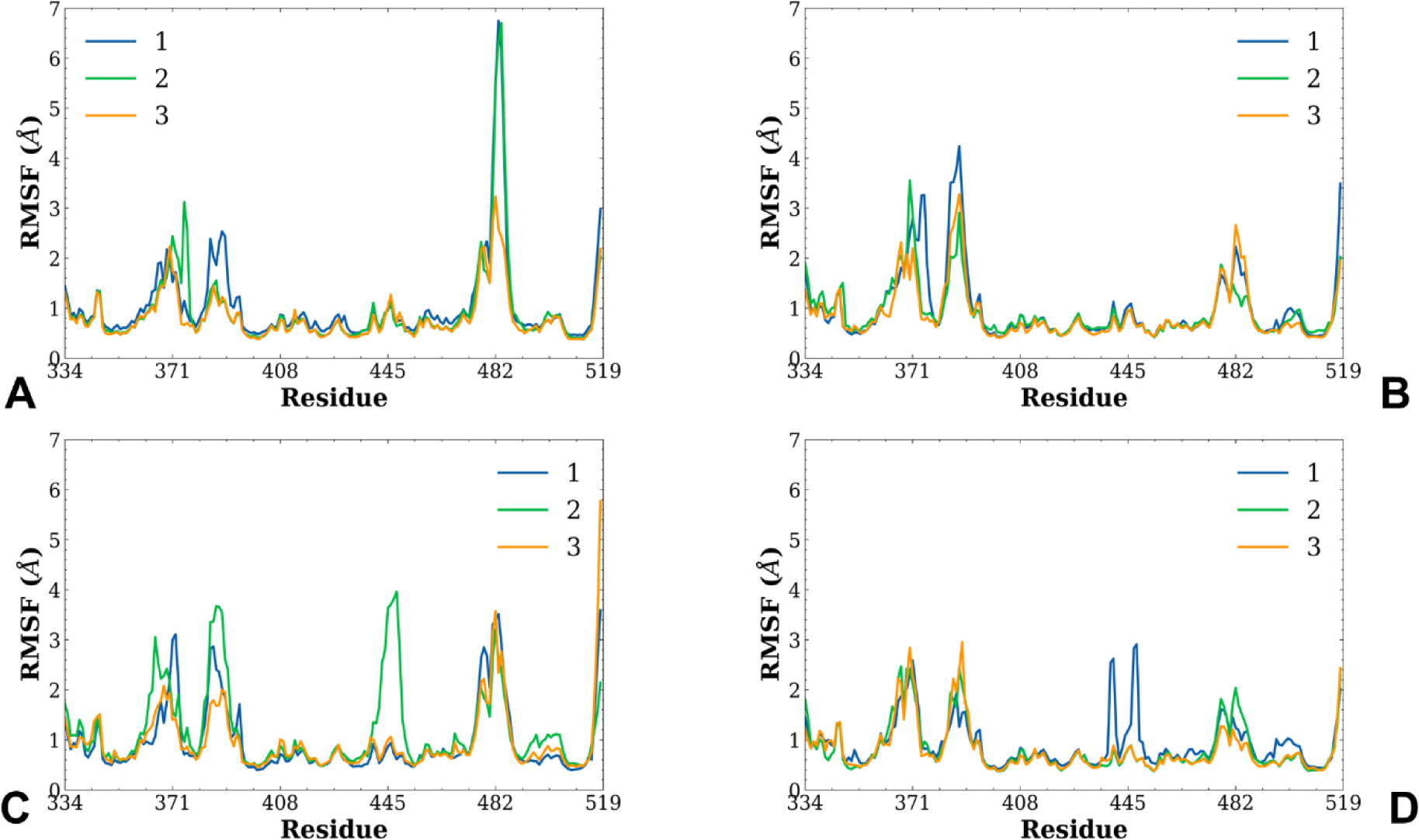
Conformational dynamics profiles obtained from three independent MD simulations of the Omicron RBD XBB.1, XBB.1.5, BQ.1 and BQ.1.1 complexes with ACE2. The RMSF profiles for the RBD residues obtained from three independent microsecond MD simulations of the Omicron XBB.1 RBD-ACE2 (A), Omicron XBB.1.5 RBD-ACE2 (B), Omicron BQ.1 RBD-ACE2 (C), and Omicron BQ.1.1 RBD-ACE2 complexes (D).

**Table 2.** Structures of the Omicron RBD-ACE2 complexes simulated in this study.

| **PDB** | **System** | **Productive run per single simulation** | **Number of simulations** |
| --- | --- | --- | --- |
| 7XB0 | Omicron RBD BA.2-ACE2 | 1μs | 3 |
| 7XWA | Omicron RBD BA.4/BA.5-ACE2 | 1μs | 3 |
| model | Omicron RBD XBB.1-ACE2 | 1μs | 3 |
| model | Omicron RBD XBB.1.5-ACE2 | 1μs | 3 |
| model | Omicron RBD BQ.1-ACE2 | 1μs | 3 |
| model | Omicron RBD BQ.1.1-ACE2 | 1μs | 3 |

The conformational flexibility of the Omicron RBD was analyzed by calculating the root mean square deviations (RMSD) (Supporting Figures S1, S2) and the root mean square fluctuations (RMSF) distribution for the RBD residues (Figure 2).

The RMSD profiles for the RBD residues showed a fast convergence of the MD trajectories, for the BA.2 and XBB.1.5 complexes where all three trajectories converged reaching a steady equilibrium state after about 350-400 ns (Supporting Figure S1). An appreciable divergence of the RMSD profiles was seen for the MD trajectories of the XBB.1 complex (Supporting Figure S1B,E) reflecting a more heterogeneous ensemble of the RBD conformations and also pointing to greater flexibility of the ACE2 enzyme. The RMSD profiles for the Omicron BQ.1.1 RBD showed convergence of the three microsecond trajectories reaching the steady state after ∼250-300 ns (Supporting Information Figure S2C), while a more significant divergence among trajectories was detected for the parental BA.5 RBD (Supporting Information Figure S2A) and particularly BQ.1 RBD (Supporting Information Figure S2B). The RMSDs for the ACE2 displayed a certain degree of variability among trajectories for BA.5, BQ.1 and BQ.1.1 variants indicating functionally significant plasticity of both binding partners in the RBD-ACE2 complexes. These preliminary findings also suggested that the RBD residues in the XBB.1 and BQ.1 variants may be more dynamic than in the very closely related XBB.1.5 and BQ.1.1 sublineages.

The RMSF profiles showed several local minima regions corresponding to the structured five-stranded antiparallel β-sheet core region (residues 350-360, 375-380, 394-403). Common to the conformational mobility profiles that displayed a similar overall shape was the presence of two RMSF peaks corresponding to the flexible RBD regions (residues 360-373 and residues 380-396) (Figure 2). The RBD regions 440-456, 470-491 and 491-505 include a number of binding interface residues. In particular, residues 491-505 forming a dominant cluster of major interaction contacts with ACE2 consistently displayed small fluctuations and featured considerable binding-induced stabilization in all complexes. Interestingly, the flexible RBD loops 440-452 and 470-491 containing many of the convergent mutation sites showed significant differences in their mobility across the Omicron variants (Figure 2).

The RMSF profiles revealed signs the greater stability for the XBB.1.5 RBD (Figure 2B) as compared to XBB.1 (Figure 2A) and parent BA.2 variant (Supporting Information Figure S1), featuring small thermal fluctuations not only in the ACE2-interacting sites but also only moderate displacements in the flexible RBD regions (residues 355-375 and 380-400) (Figure 2A,B). The stable RBD core regions (residues 400-420, 430-450) exhibited even smaller fluctuations in the XBB.1.5 complex (Figure 2A,B), suggesting the increased RBD stability for these variants which may be a relevant contributing factor to the stronger ACE2 binding experimentally observed for XBB.1.5 subvariant.^54,55^ The RMSF profile for XBB.1.5 is characterized by several local minima corresponding to the rigid RBD core, the ACE2 interfacial sites (residues 485-505) (Figure 2B). Importantly, the thermal movements of the intrinsically more flexible RBM region were markedly suppressed in the XBB.1.5 variant (Figure 2B). The RBM tip in the XBB.1.5 RBD-ACE2 complex is maintained in a stable folded conformation that can be described as a hook-like folded RBD tip and is similar to the crystallographic conformations. Interestingly, in the XBB.1 RBD, the RBD tip is more flexible and samples a more dynamic state in which the RBD tip circulates between a variety of partly disordered conformations (Figure 2A).

By constructing mutational models of BQ.1 and BQ.1.1 RBD-ACE2 complexes using the crystal structure of the BA.4/BA.5 complex and performing microsecond atomistic MD simulations, we observed several interesting patterns in the distributions. The RMSF profiles for the BQ.1 RBD residue showed markedly greater fluctuations (Figure 2C) as compared to closely related BQ.1.1 RBD (Figure 2D). In particular, we noticed the reduced mobility in the RBM region and partly suppressed motions of intrinsically flexible RBM tip residues 475-486 in the BQ.1.1 (Figure 2D). Interestingly, convergent mutation sites R346T, K444T and N460K belong to fairly exposed flexible regions, but these positions experienced only very moderate fluctuations in BQ.1.1 RBD-ACE2 complex. One of the microsecond MD trajectories of the BQ.1.1 RBD-ACE2 complex showed some divergence in the RMSF values showing larger fluctuations of the convergent mutational positions K444, V445, G446 (Figure 2D). The key Omicron mutational positions Q493R, Q498R, N501Y and Y505H that participate in the critical ACE2 binding interface region are highly constrained by their strong interactions with the receptor and experience only very minor fluctuations across all the studied variants. Overall, the important difference in the dynamic signatures between Omicron variants was a noticeable reduction in the mobility of the intrinsically flexible RBM loop for XBB.1.5 and BQ.1.1 subvariants, while this region is characterized by greater conformational variability and lack of the well-defined folded RBM tip in the XBB.1 and BQ.1 variants (Figure 2).

The RMSF analysis of the ACE2 residues showed similar profiles across all the examined variants (Figure 3). The highly stable ACE2 residues corresponding to the rigid core and the binding interface positions centered around K353 and H34 (Figure 3). The key ACE2 binding motifs correspond to an alpha-helix (residues 24-31) and a beta-sheet (residue 350-356) that display moderate RMSF values in all complexes. The important polar /charged residue interactions at the interface are formed with ACE2 residues D30, K31, H34 and E35 that all display exceedingly small thermal fluctuations. Other ACE2 residues Q24, M82, Y83, D38, Y41, N330, K353 that anchor distinct parts of the RBD-ACE2 binding interface remained stable in all MD trajectories (Figure 3). Notably, the conformational dynamics profiles revealed signs of the increased ACE2 heterogeneity in the flexible loop (residues 340-350) located away from the binding interface, and this mobility is particularly increased in the XBB.1.5 and BQ.1.1 RBD-ACE2 complexes (Figure 3B,D).

**Figure 3.**
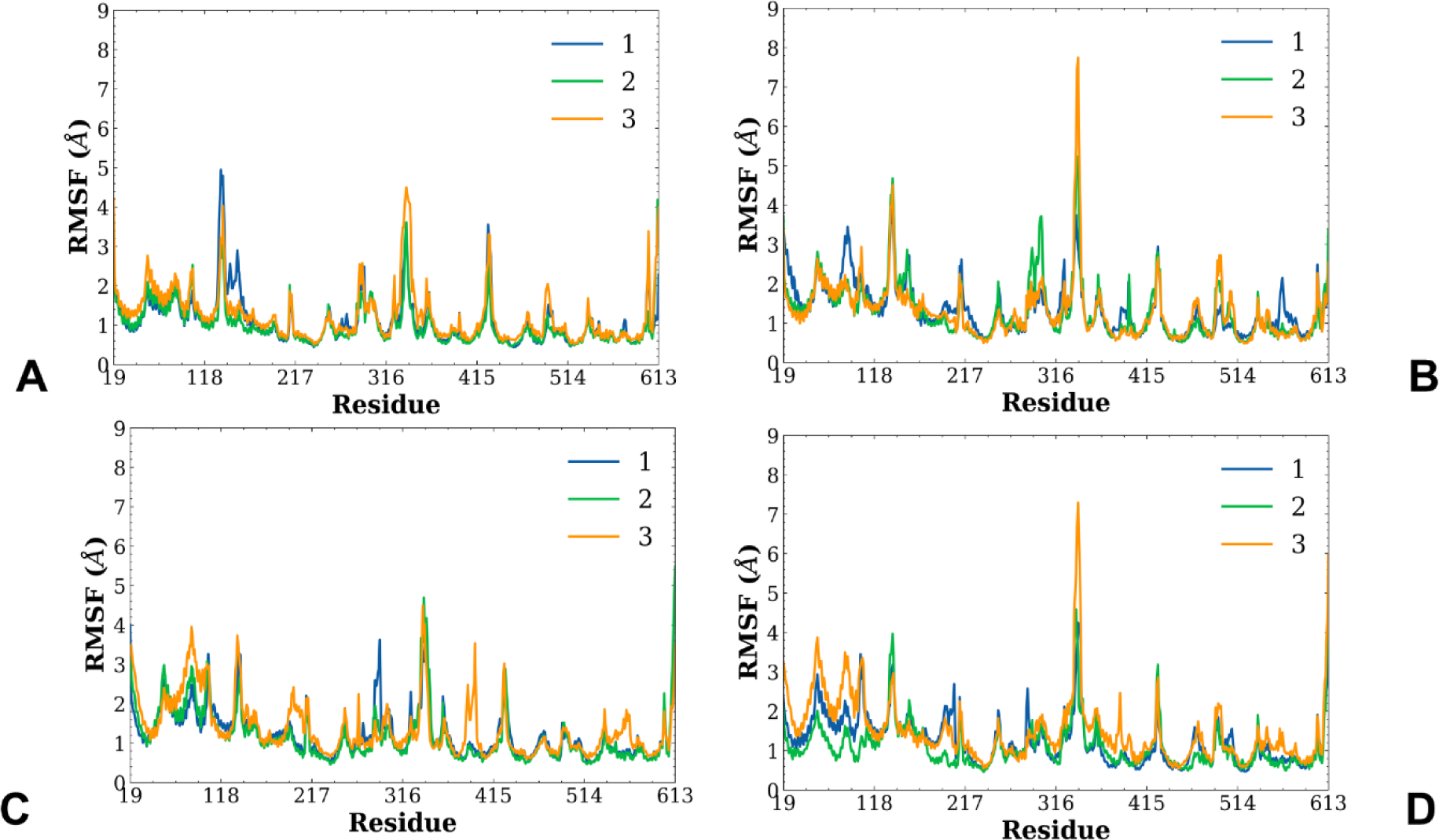
Conformational dynamics profiles of the ACE2 residues obtained from three independent MD simulations of the Omicron RBD XBB.1, XBB.1.5, BQ.1 and BQ.1.1 complexes with ACE2. The RMSF profiles for the ACE2 backbone residues obtained from three independent microsecond MD simulations of the Omicron XBB.1 RBD-ACE2 (A), Omicron XBB.1.5 RBD-ACE2 (B), Omicron BQ.1 RBD-ACE2 (C), and Omicron BQ.1.1 RBD-ACE2 complexes (D).

To further examine the character of dynamic couplings and quantify correlations between motions of the RBD regions we performed the dynamic cross correlation (DCC) residue analysis and reported the DCC maps for the Omicron RBD-ACE2 complexes (Figure 4). The DCC maps demonstrated subtle differences in the dynamic couplings (Figure 4). For XBB.1 and XBB.1.5 RBD complexes, we observed strong positive correlations among residues 395-475 (Figure 4A,B).

**Figure 4.**
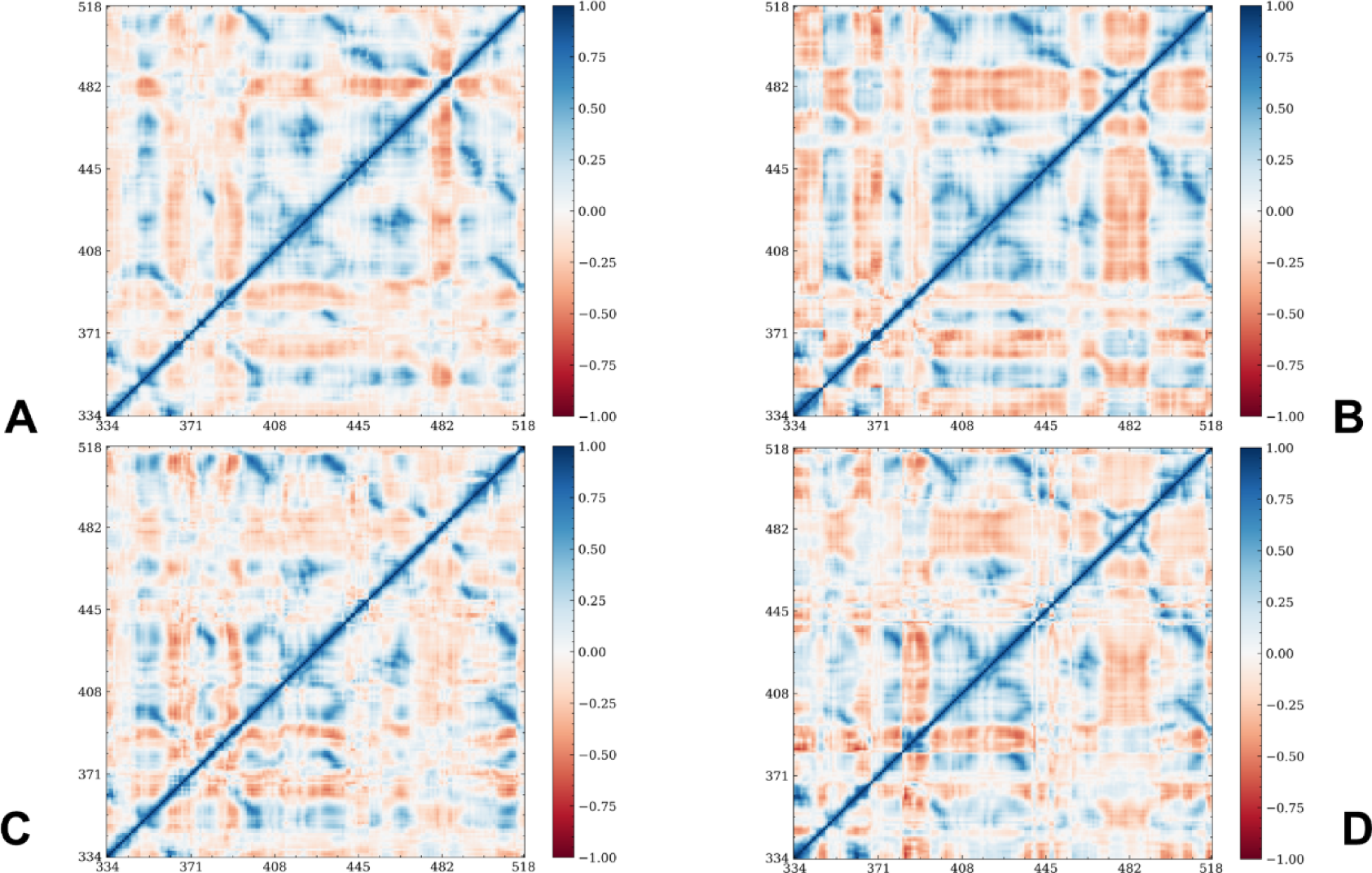
The DCC for the RBD residues in the Omicron XBB.1 RBD-ACE2 (A), Omicron XBB.1.5 RBD-ACE2 (B), Omicron BQ.1 RBD-ACE2 (C), and Omicron BQ.1.1 RBD-ACE2 complexes (D).

The movements of these RBD residues are anti-correlated with the displacements of the RBM tip (residues 475-487). This reflects a more stable RBD in the XBB.1.5 complex yielding strong dynamic couplings between the RBD core and more flexible regions of the binding interface. Hence, conformational dynamics of the XBB.1.5 RBD exhibited signs of long-range allosteric couplings between the RBD core elements and flexible loops that could collectively determine the RBD response to binding (Figure 4A,B). A different pattern of dynamic couplings was observed for BQ.1 (Figure 4C) and BQ.1.1 (Figure 4D). For BQ variants, positive dynamic correlations were seen for residues 335-375 and 396-455 in the RBD, while movements of the flexible loop (residues 375-395) are anticorrelated with the RBD core. Common to XBB.1.5 and BQ.1.1 RBDs is the presence of negative cross-correlations between motions of the RBM tip (residues 475-487) and other RBD regions (residues 400-470, 490-520) (Figure 4), where the extent of anticorrelation is stronger for XBB.1.5 RBD (Figure 4B). The key differences in the conformational dynamics of the Omicron RBD-ACE2 complexes were further exemplified by the DCC maps pointing to the reduced mobility of the RBM flexible loops in XBB.1.5 and BQ.1.1 and the enhanced long-range dynamic couplings between RBD regions in these complexes (Figure 4).

To summarize, microsecond MD simulations of the Omicron RBD-ACE2 complexes presented convincing evidence of distinct dynamic patterns in the structurally similar Omicron RBD-ACE2 complexes. Consistent with the experimental data, our results showed that XBB.1.5 and BQ.1.1 mutations may induce the increased stabilization of the RBD in the complex with ACE2 which may be linked with the greater RBD stability and enhanced binding as compared to the other variants.

### Dimensionality Reduction Analysis Unveils Differences in Conformational Heterogeneity of the Omicron RBD-ACE2 Complexes

While the analysis of MD simulations and functional movements provided important insights into the underlying conformational landscape the high dimensionality of the data sets produced by simulations often hinders salient dynamic signatures associated with the mechanisms of allosteric transitions. Here, to facilitate the conformational landscape analysis we employed a dimensionality reduction method to project the results of MD simulations into low dimensional space.^106-109^ Given a time-series of molecular coordinates provided by the MD trajectories, tICA aims to reduce the dimensionality of the trajectories and to identify hidden key structural changes. The low-dimensionality projection of the MD ensemble in different microsecond trajectories highlighted a consistent occupancy of the dominant conformational region for BA.2 and (Figure 5A) its descendants XBB.1 (Figure 5B) and XBB.1.5 variants (Figure C). In striking contrast, a larger conformational space is available for BQ.1 (Figure 5E) and BQ.1.1 (Figure 5F). Interestingly, the conformational space of the parental BA.4/BA.5 variant remains narrow and similar to BA.2. This analysis showed that a small number of mutational changes acquired in BQ.1 and BQ.1.1 variants due to accumulation of convergent mutations in R346, K444, L452 and N460 positions could lead to a significant enhancement in conformational variability and incur redistributions in the populations of different states (Figure 5). The broader area of the conformational space for BQ.1 reflected the greater mobility of BQ.1 RBD, while BQ.1.1 may sample some additional regions of the conformational space. Notably, the dominant conformational region in the low-dimensional space is shared among all variants and it corresponds to the crystallographic conformations observed in the parental BA.2 and BA.4/BA.5 variants. Moreover, the dominant low-dimensional region is associated with “hook-like” folded conformation of the RBM tip which is observed in the crystallographic conformations. The heterogeneity of the dynamic states revealed in the low-dimensional projection space for BQ.1 and BQ.1.1 (Figure 5E,F) suggested that convergent mutations may induce unique dynamic signatures of the conformational ensembles for these variants.

**Figure 5.**
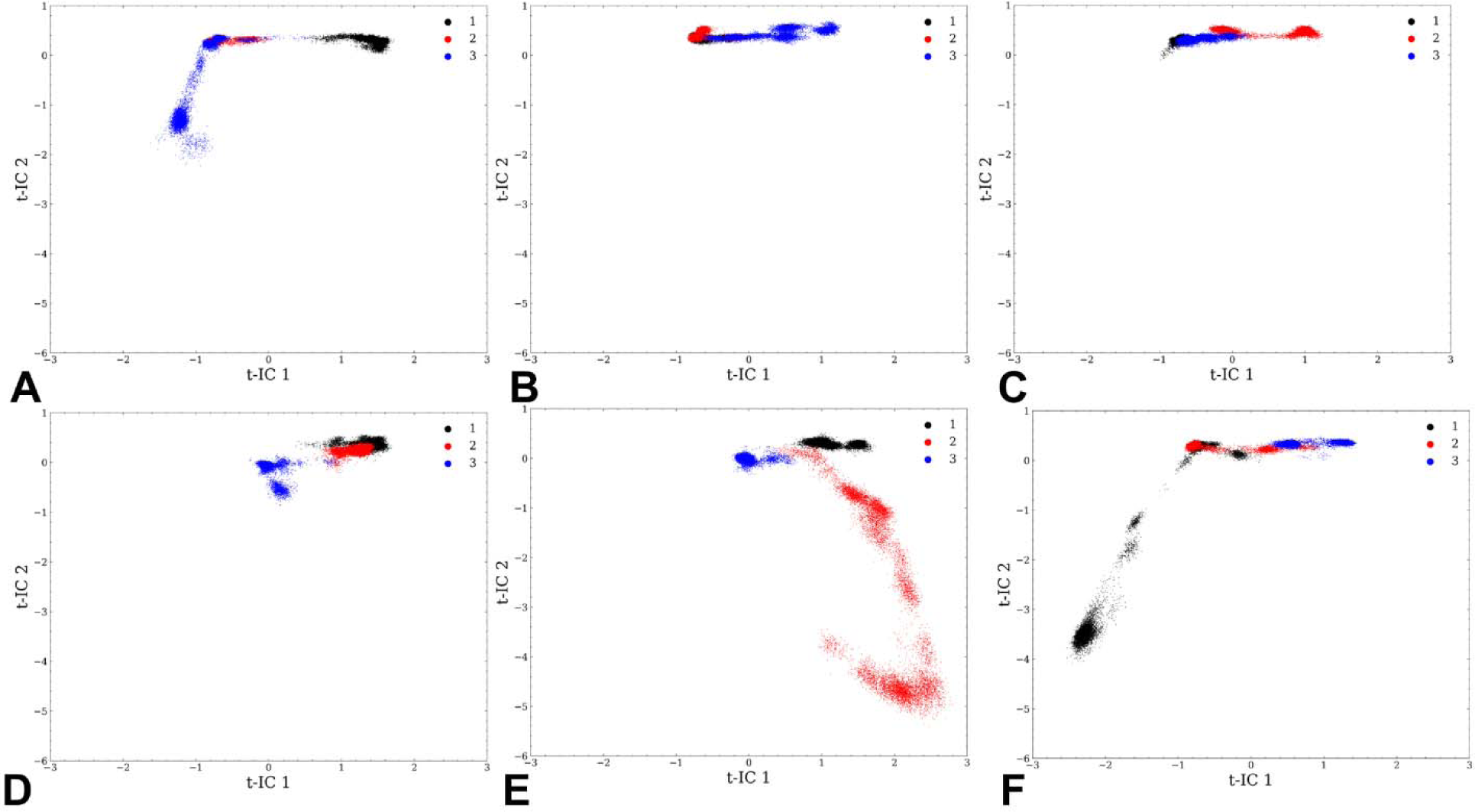
The t-ICA low-dimensional projection of the conformational space sampled in three independent microsecond MD simulations for the Omicron XBB.1 RBD-ACE2 (A), Omicron XBB.1.5 RBD-ACE2 (B), Omicron BQ.1 RBD-ACE2 (C), and Omicron BQ.1.1 RBD-ACE2 complexes (D).

A common and powerful approach to probe conformational landscapes is to combine MD simulations with MSM analysis to estimate probabilities for protein transition among different macrostates. To build effective MSM, it is important for a dimensionality reduction method to retain information about how proteins transition among these macrostates. To evaluate the retaining of such information, we analyzed the relaxation timescales in MSM, also referred to as implied timescales. The relaxation timescale can be interpreted as the time needed for a system to change its state. To cluster different conformations into metastable states, *k*-means clustering method was used to build clusters with the mean RMSD within cluster smaller than 1 Å. In the t-ICA reduced 2D space, k-means clustering method was applied to partition the 2D data into 20 states. The implied time scales were calculated with lag times ranging from 0.1 to 5 ns. The top 15 timescales are shown (Figure 6A). The trend of implied relaxation time scale revealed that the estimated time scale converged after ∼3 ns, which was chosen as the lag time in the construction of MSM. Based on the gap of time scales, the number of macrostates was set to seven. The low-dimensional representation with 2 components (t-IC) showed some overlapping and the increased density of states in the dominant region sampled in all examined Omicron RBD-ACE2 systems (Figure 6B). The high density regions are predominantly sampled in BA.2, XBB.1 and XBB.1.5 variants (Figure 6B) and these regions are mainly associated with macrostates 3 and 4 (Figure 6C). The less populated regions are sampled in the BQ.1 and BQ.1.1 variants where macrostates 1,2,5, 6 and 7 contribute to the overall distribution of states (Figure 6B,C). To visualize the distribution density of all simulations, a hexagon binning was employed where each hexagon represents a small region, and the color bar shows the frequency of this region (Figure 6D). The density plot revealed a clearly dominant area in the low-dimensional space sampled by all variants. Importantly this region is exclusively sampled by XBB.1.5 variant, suggesting considerable stabilization of this complex. The least dense area of the distribution (“right leg”) appeared mostly in a highly dynamic BQ.1 variant. In general, the low-dimensional projection analysis emphasized the key differences in conformational dynamics of the Omicron variants, where XBB.1 and XBB.1.5 RBDs tend to occupy a more limited area of the conformational space corresponding to the crystallographic conformation of the parental BA.2. In contrast, convergent mutations in BQ.1 and BQ.1.1 may induce conformational heterogeneity which is particularly apparent in the BQ.1 variant with the most flexible RBD structure among studied systems.

**Figure 6.**
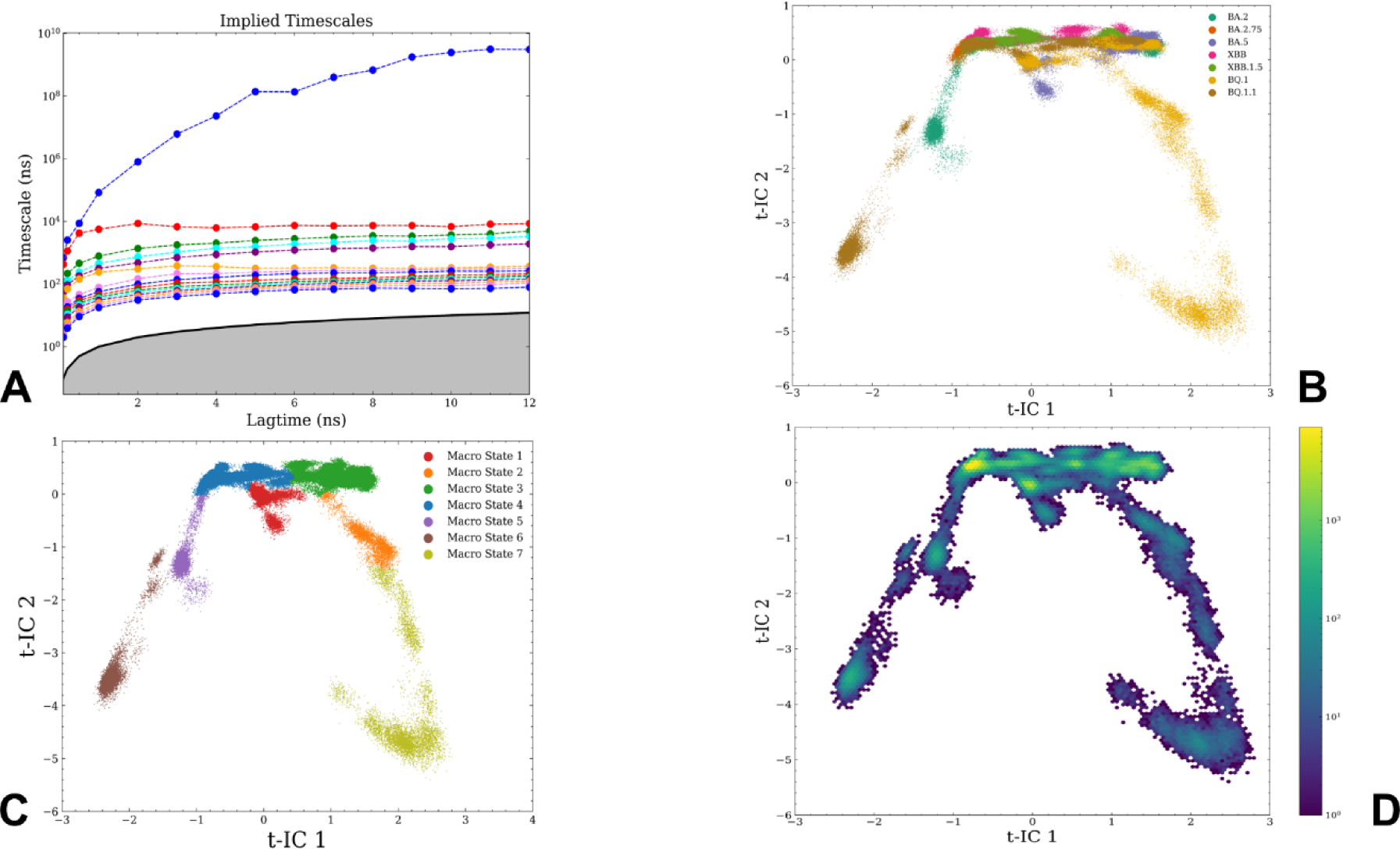
An overview of the MSM analysis of the conformational landscapes for Omicron RBD-ACE2 complexes. (A) The implied time scales were calculated with lag times ranging from 0.1 to 5 ns. The top 15 timescales are shown. The estimated time scale converged after ∼3 ns, which was chosen as the lag time in the construction of MSM. (B) The low-dimensional density representation with 2 components (t-IC) is constructed based on the MSM analysis. The distribution densities are colored based on the system and are shown for BA.2 (in emerald green), BA.2.75 (burnt Sienna), BA.5 (purple), XBB.1 (pink), XBB.1.5 (forest green), BQ.1 (earthy yellow) and BQ.1.1 variants (brown). (C) The low-dimensional density representations with 2 components (t-IC) are shown for macrostate 1 (red), macrostate 2 (orange), macrostate 3 (green), macrostate 4 (blue), macrostate 5 (purple), macrostate 6 (brown), and macrostate 7 (lime zest). (D) The distribution density of all simulations is visualized using a hexagon bin plot. Each hexagon represents a small region, and the color bar shows the frequency of this region.

After the partition of the MSM macrostates to macrostates was determined (Supporting Information Figure S3), the stationary distribution and transition probabilities were calculated based on the constructed MSM. We determined the distributions of the Omicron RBD-ACE2 complexes in different macrostates which reveal the unique dynamic signatures and populations in each complex. The transition probabilities are determined among different macrostates for all systems, with 3 ns lag time. The high percentage of self-conserved probability shows the stability of macrostates. We first compared the distributions and transitional maps between parental BA.2 (Supporting Information Figure S4A) and XBB.1/XBB.1.5 subvariants (Figure 7A,B). For the BA.2 variant, we found that macrostates 3,4,5 dominate the distribution with the macrostate 4 contributing ∼51 % to the conformational population (Figure 7A). The transitional map for BA.2 showed that the macrostates could have high kinetic barriers for transition to the other states (Supporting Information Figure S4B). For example, for the macrostate 4 there is an overall probability of less than 0.5 % to shift to two macrostates 3 and 5.

**Figure 7.**
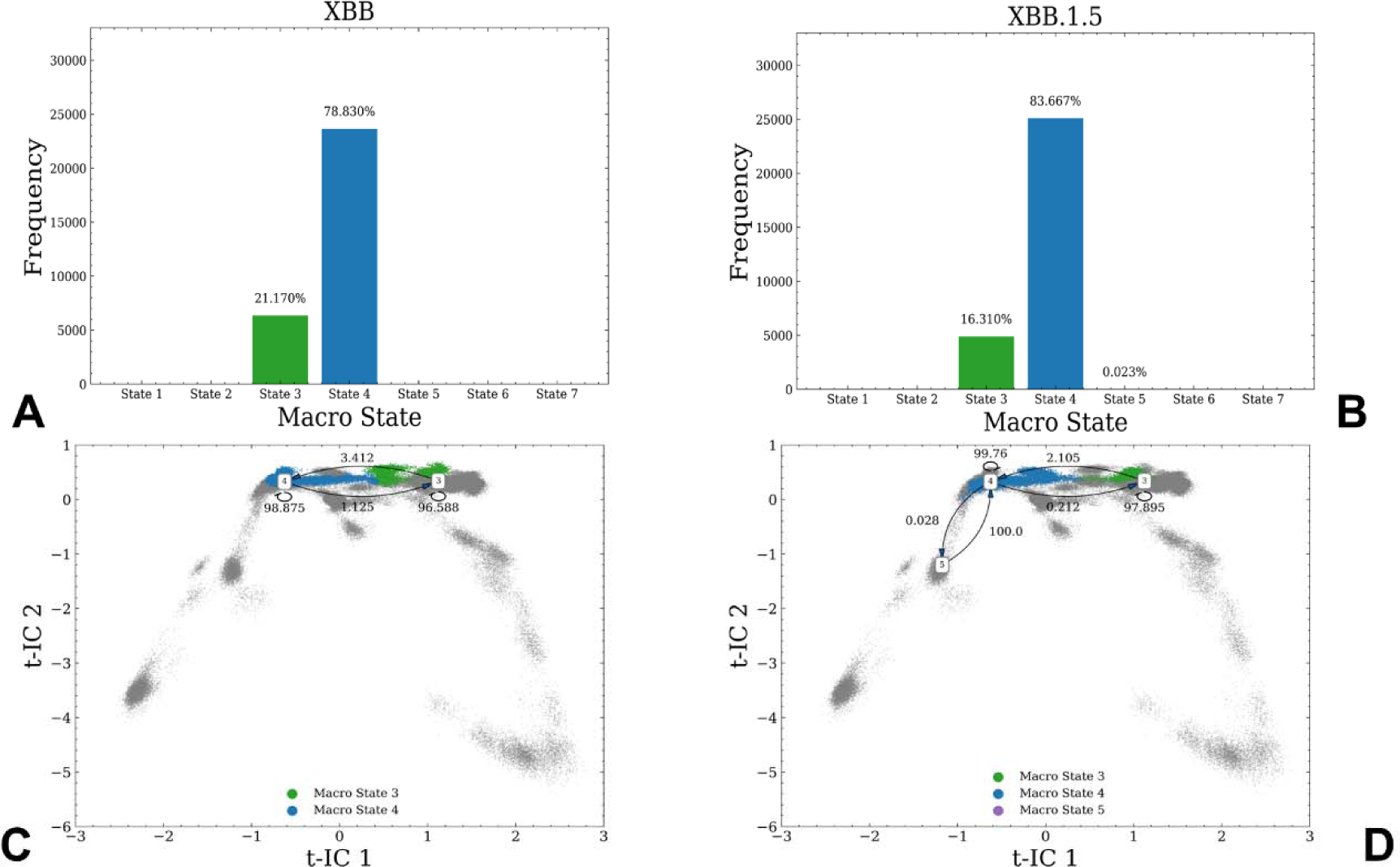
MSM analysis of the conformational landscape for XBB.1 and XBB.1.5 RBD-ACE2 complexes. The frequency distribution of occupying different macrostates in XBB.1 (A) and XBB.1.5 (B). The transition probability maps among different macrostates with 3 ns lag time for XBB.1 (C) and XBB.1.5 variant (D). The high percentage of self-conserved probability shows the stability of macrostates.

At the same time, there is almost a 100% probability that minor macrostate 6 would interconvert to the macrostate 5 but close to zero probability of the reverse transition (Supporting Information Figure S4B). A comparative MSM analysis for XBB.1 and XBB.1.5 sublineages of BA.2 showed a reduction in the number of macrostates, where macrostates 3 and 4 begin to dominate the conformational distribution for XBB.1 (Figure 7A,C) and XBB.1.5 RBD (Figure 7B,D). Moreover, the population of the macrostate 4 increased in the XBB.1.5 RBD to 86% from 75% in the XBB.1 variant (Figure 7B).

Structural analysis of the obtained macrostates showed that the differences between macrostates 3 and 4 are confined to the flexible RBM region (residues 470-491) that harbors several Omicron XBB.1/XBB.1.5 mutational sites and convergent Omicron sites F486P (F486S), F490 (F490S) (Figure 8). The MSM analysis revealed that dominant macrostate 4 featured well-ordered and stable “hook-like” RBM tip conformation. At the same time, macrostates 4 and 5 in BA.2 have similar ordered RBM tip region but differ in the conformation of the flexible loop 440-452 (Figure 8). In macrostate 4 this loop assumes the crystallographic conformation of the BA.2 RBD, while this loop becomes more disordered in macrostate 5 (Figure 8). Hence, the major structural differences between major macrostates are mostly confined to the order-disorder transitions in the flexible loop regions 440-452 and 470-491. Interestingly, these regions harbor a number of convergent mutational sites including K444, V445, G446, N450, L452 positions.

**Figure 8.**
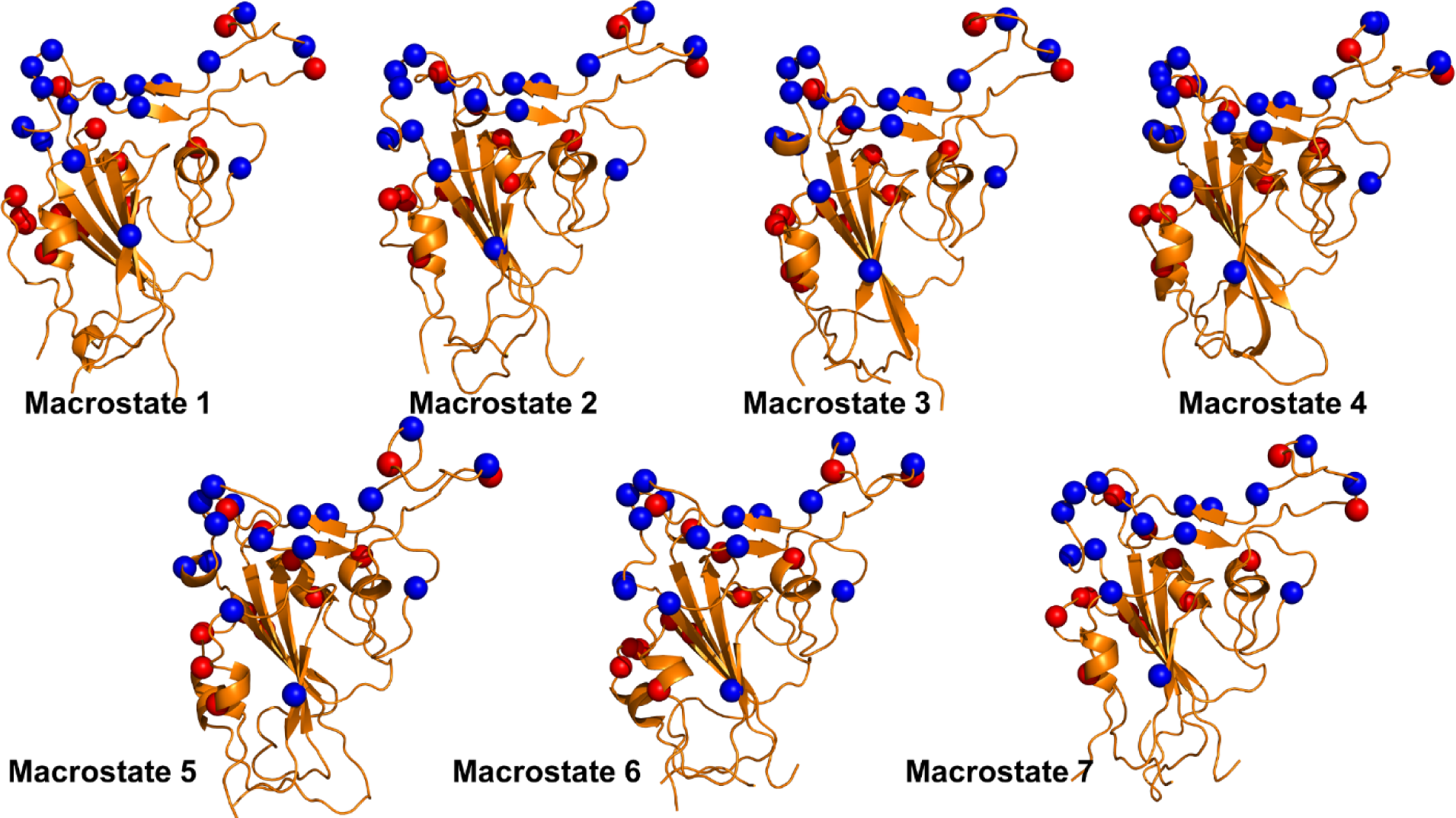
Structural analysis of macrostates determined in the MSM analysis. To perform a comparative structural analysis of obtained macrostates, the assignments of all points are examined to determine which point belongs to which macrostate. By calculating the centroid of the conformations that belong to each macrostate, the closest to the centroid existing frame is determined and considered as a representative of a given macrostate. The macrostates are annotated and are shown in orange-colored ribbons. The convergent mutation sites are highlighted in blue spheres. The analysis shows differences between conformations that are particularly exemplified by changes in the flexible loops 440-452 and 470-491.

The transition maps for XBB.1 (Figure 7C) and XBB.1.5 (Figure 7D) unveiled a small transitional probability to interconvert from macrostate 4 to macrostate 5, indicating the increasing dominance of macrostate 4. The MSM analysis also revealed that dominant macrostate 4 featured well-ordered and stable “hook-like” conformation of the RBM tip centered on F486P position in XBB.1.5 (Figure 8). This state becomes less favorable in XBB.1 while the contribution of macrostate 3 with a partly disordered conformation of the RBM tip increased. Based on the structural comparison of the macrostates, it becomes apparent that a combination of F486S and F490S mutations in XBB.1 may induce the increased mobility of the flexible RBM tip. On the other hand, the effect of F486P mutation in XBB.1.5 variant can be manifested in partial restoration and stabilization of the interfacial contacts mediated by F486P in this region. This analysis provides a more granular landscape-derived rationale for the experimental data showing that XBB.1.5 is equally immune evasive as XBB.1 but may have growth advantage by virtue of the higher ACE2 binding as F486P in the XBB.1.5 subvariant can restore most of the favorable hydrophobic contacts. According to DMS experiments among the most common F486 mutations (F486V/I/S/L/A/P), F486P imposes the lowest cost in RBD affinity loss and has the largest increase in RBD expression.^131,132^ F486V (BA.4/BA.5), F486I, F486S (XBB.1), F486P(XBB.1.5) also represents a convergent evolutionary hotspot shared by the recent wave of Omicron subvariants and is one of the major hotspots for escaping neutralization by antibodies.^133^

A comparison of the macrostate distributions and transition maps for the parental BA.5 (Supporting Information Figure S4B) and BQ.1/BQ.1.1 variants (Figure 9) revealed a vastly different pattern even though the structures of all Omicron RBD-ACE2 complexes are similar. Interestingly, for the parental BA.5 variant, we observed populations of macrostate 3 and macrostate 1, where structurally macrostate 1 is more similar to the experimental structure and has a well-ordered RBM tip conformation (Supporting Information Figure S4B). Hence, the increased mobility of the BA.5 RBD is manifested in a more populated state with the very flexible RBM region (residues 470-491). Interestingly, the distribution is broadened in the BQ.1 variant showing significant contribution of macrostates 1 (34%), 2(11%), 3 (35%) and 7 (20%) (Figure 9A,C). Structural analysis showed that macrostates 3 and 7 are similar in their partly distorted RBM tip conformation abrogating “hook-ordered” conformation, but these macrostates are different in the conformation of the loop 440-452 (Figure 8). The macrostate 1 has a well-ordered RBM tip similar to the crystallographic conformation of BA.5 but featured conformational differences distributed over the flexible loops in the RBD (Figure 8).

**Figure 9.**
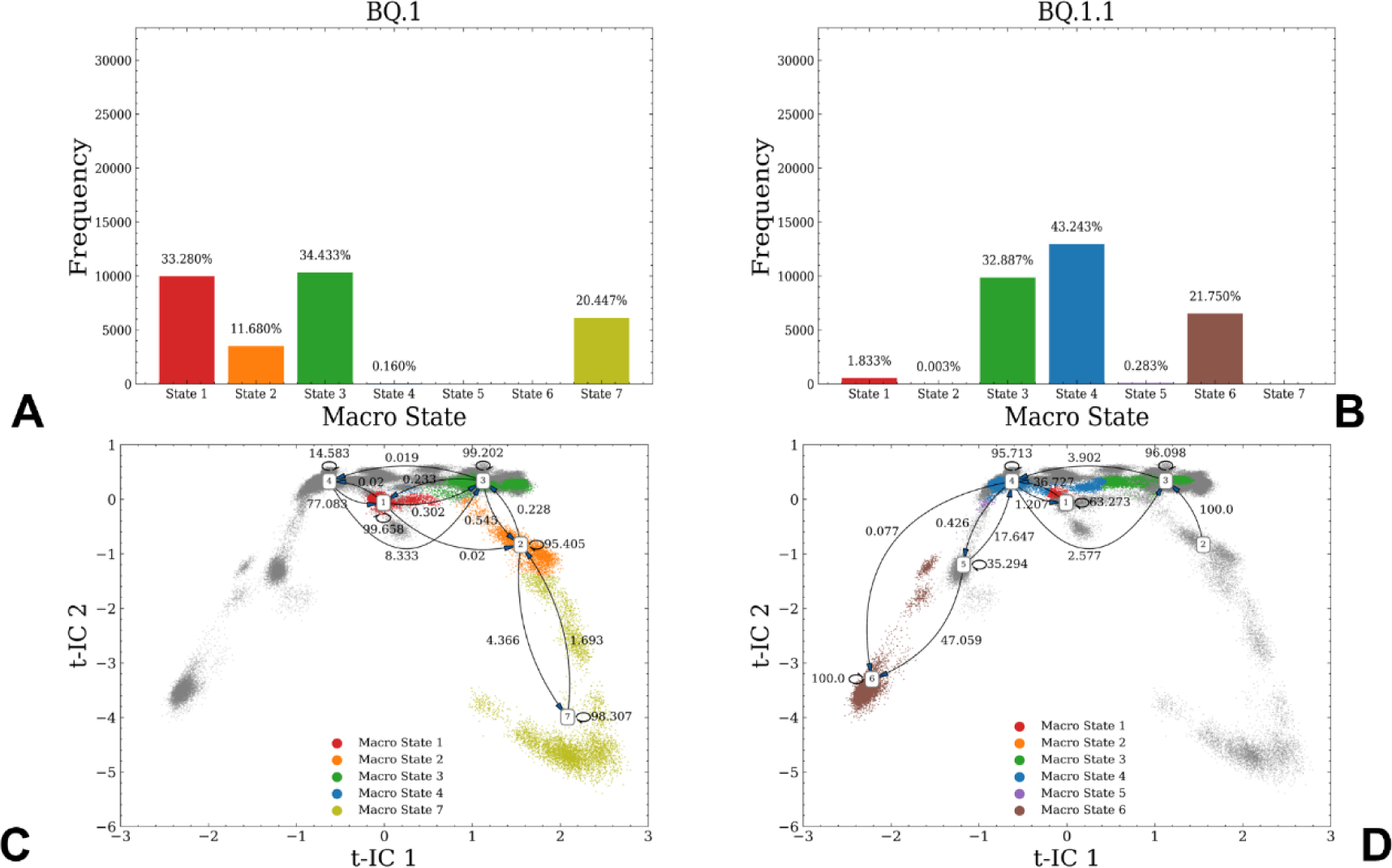
MSM analysis of the conformational landscape for BQ.1 and BQ.1.1 RBD-ACE2 complexes. The frequency distribution of occupying different macrostates in BQ.1 (A) and BQ.1.1 (B). The transition probability maps among different macrostates with 3 ns lag time for BQ.1 (C) and BQ.1.1 variant (D). The high percentage of self-conserved probability shows the stability of macrostates.

The transitional maps showed that macrostates 3,1 and 4 occupy neighboring areas of conformational space (Figure 9). Interestingly, the distribution of macrostates is altered in the BQ.1 variant (Figure 9A,C), as macrostate 1 begins to dominate the population while the contribution of macrostate 4 is markedly diminished. The transition map for BQ.1 showed that macrostate 4 can convert to macrostate 1 with 77% probability while the reversed transition is highly unfavorable (Figure 9C,D). Macrostate 3 is also connected with macrostates 2 and 7 in the BQ.1 variant. Overall, the dominant population of macrostates for BQ.1 is characterized by distorted and partly disordered RBM tip conformation featured in macrostates 3 and 7 and enhanced flexibility of the RBD loops in macrostate 1. These findings detailed and provided further support to the observed conformational plasticity of the BQ.1 variant that is mediated by convergent mutations and allows for the enhanced immune escape potential.

Remarkably, the MSM analysis revealed that the distribution of macrostates is markedly changed in a closely related BQ.1.1 variant (Figure 9B,D). We observed the population shift to macrostates 3,4 and 6 dominating the conformational distribution, with macrostate 4 contributing 43%, macrostate 3 contributing 33 % and macrostate 6 providing 22 % of the total population (Figure 9B,D). While macrostate 3 has a partially disordered RBM tip, macrostate 6 displayed alterations in the flexible loop 440-452 as compared to the crystallographic conformation of BA.5 RBD (Figure 9B,D). The transition map showed that macrostate 5 could interconvert to macrostate 4 with ∼17% probability but the interconversion between states 4 and 3 is less probable (with only 3.9% from state 3 to 4 and 2.5 % of the reversed transition) (Figure 9D). At the same time, macrostate 2 could readily interconvert to macrostate 3 with a 100% probability and macrostate 6 can convert to macrostate 5 with a 47% probability (Figure 9D).

Although the distribution of macrostates in BQ.1.1 becomes more similar to that of XBB.1.5 variant, revealing the critical contribution of macrostates 3 and 4, the population of states is significantly broader than in XBB.1.5 (Figure 9C,D). In addition to the most dominant macrostate 4, the combined contribution of macrostates 3 and 6 is equally significant. These macrostates feature distinct variations of the flexible RBD loops 440-452 and 470-491. As a result, BQ.1.1 RBD may exist in a dynamic equilibrium between the crystallographic conformation seen in the parental BA.5 RBD and more dynamic states with highly variable RBM loops (Figure 9C,D). These results demonstrated that convergent mutations that are characteristic BQ.1 and BQ.1.1 variants may significantly change the dynamic equilibrium and modulate the increased flexibility of the RBD as compared to the more stable XBB.1.5 variant.

Overall, the MSM analysis provided in-depth characterization of conformational macrostates, revealing the unique and distinct dynamic signatures of structurally similar Omicron RBD-ACE2 complexes. One of key findings of this analysis revealed a strong preference of XBB.1.5 for a single macrostate associated with the ordered RBM conformation due to the restoration of F486P contacts with ACE2 inducing curtailed mobility in this region. Our results suggest that this dynamic signature may allow XBB.1.5 to maintain immune evasion while protecting the RBD stability and binding. In contrast, the dominant macrostates in XBB.1 are characterized by highly flexible RBM conformation induced by F486S mutation and weaker ACE2 contacts which may explain the growth advantages of XBB.1.5 variant. Another key finding is the emergence of several dominant macrostates in the BQ.1.1 variant including conformations with highly flexible loop 440-452. Based on our results, BQ.1.1 variant mutations (R346T, K444T and N460K) can increase the mobility in the loop harboring convergent mutational sites K444T, V445, G446. Notably, BQ.1.1 bears all five recent convergent mutations: R346T, K444T, L452R, N460K, F486V. The increased adaptability of the 440-452 loop may provide means for the enhanced immune evasion of the BQ.1.1 variant, further emphasizing the role of convergent mutations in modulating flexibility of the mobile regions.

In general, MSM analysis consistently showed that major macrostates in the Omicron variants may differ through modulation of mobility in the intrinsically flexible RBD loops 470-491 and 440-452 involved in the interfacial contacts and containing sites of convergent mutations K444, V445, G446, N450, L452 and F486. The variant-specific changes of conformational mobility in these functional loops may be partly modulated and altered through accumulation of various convergent mutations providing evolutionary path for improved immune escape without significant cost on the ACE2 binding. The immune pressure on the RBD that promotes convergent evolution may be localized in these functional loops responsible for dynamic differences between variants. We argue that through this interplay of dynamic changes modulated by convergent mutations BA.2/XBB.1/XBB.1.5 or BA.5/BQ.1/BQ.1.1 subvariants can enhance antibody escape.

### Perturbation Response Scanning Reveals Complementary Role of Convergent Mutations as Allosteric Effector and Sensor Centers

Patterns of convergence are often offered as compelling evidence of adaptation and recent studies indicated that allosteric interactions in proteins not only dictate long-range couplings and signal transmission patterns but are also essential in the evolutionary adaptation of proteins.^134,135^ It has been recognized that allosteric mechanisms regulate emergence and contribution of different conformational states and shape up functional landscapes, enabling evolutionary adaptation of new functions through controlled modulation of conformational plasticity.^135^ We hypothesized that sites of convergent mutations target specific structural hotspot of adaptation that may be organized through engagement to the pre-existing allosteric network. In this model, convergent mutations may work in concert with each other to potentiate functional protein adaptation to binding and immune escape via modulation of conformational flexibility in the key regions. According to the MSM analysis, we suggest that specific modulation of mobility in the two flexible regions (residues 440-452 and 470-491) may be orchestrated through allosteric interaction network that couples sites convergent mutations.

To explore potential allosteric roles of the convergent mutations in RBD and characterize how Omicron variants can exploit these sites to optimize functional adaptation, we employed perturbation profiling methodology to interrogate the inter-residue couplings and particularly probe the role of convergent mutations in mediating long-range allosteric effects in the Omicron RBD-ACE2 structures. The PRS method^119-121^ was used to examine the allosteric roles of the RBD residues using the conformational ensembles of the XBB.1/XBB.1.5 and BQ.1/BQ.1.1 RBD-ACE2 complexes (Figure 10). The effector profiles measured the allosteric capacity of residue *i* to influence dynamics changes in all other residues, with the local maxima usually attributed to allosteric centers that may control propagation of perturbations over long distance and allosteric communications in the system. At the same time, the transmission capacity of residue *j* to perturbations can be evaluated from sensor distributions in which major peaks are often aligned with flexible regions experiencing large functional changes.

**Figure 10.**
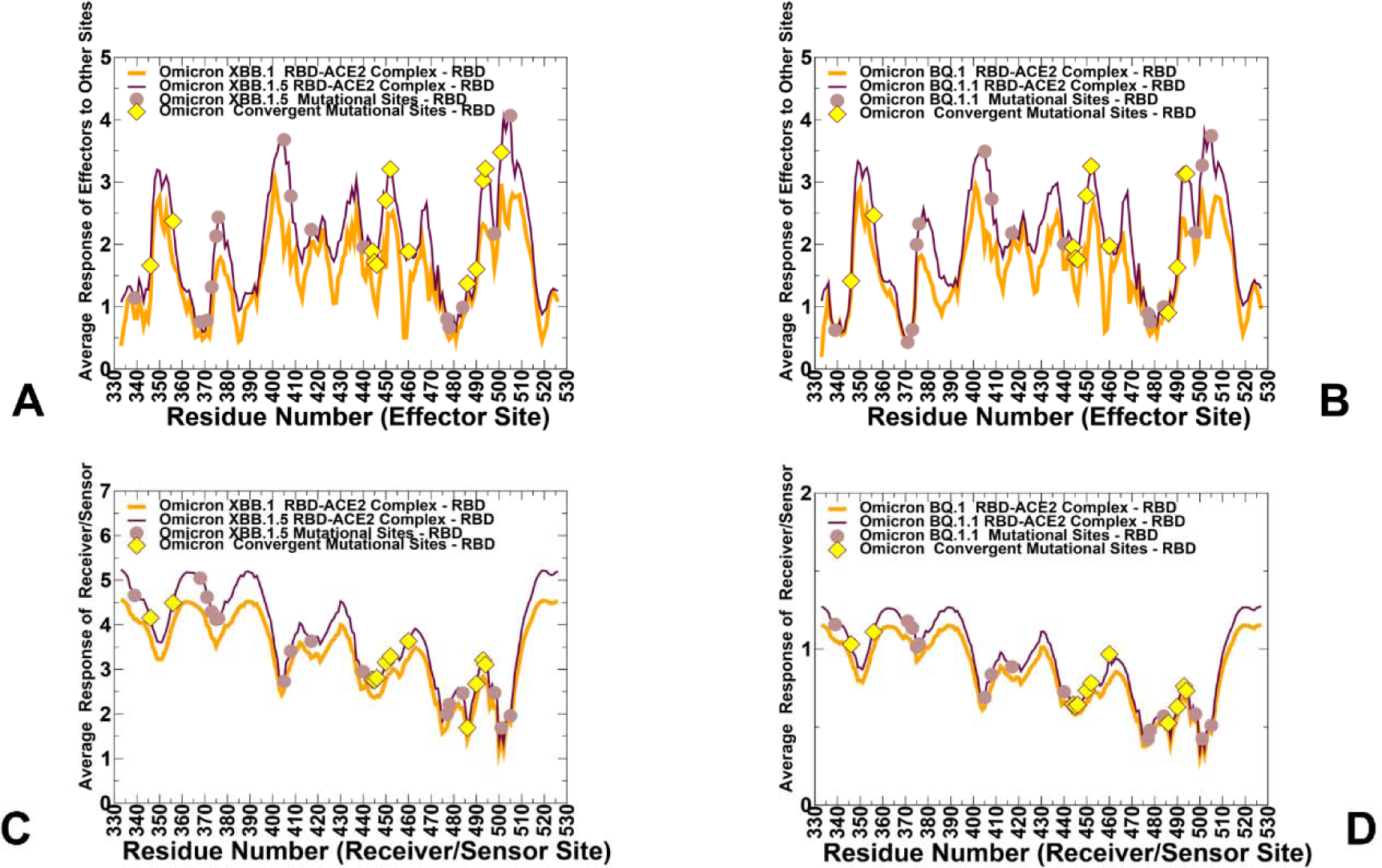
The PRS analysis of the SARS-CoV-2 Omicron RBD-ACE2 complexes. (A) The PRS effector profiles for the XBB.1 RBD (in orange lines) and XBB.1.5 RBD (in maroon lines). (B) The PRS effector profiles for the BQ.1 RBD (in orange lines) and BQ.1.1 RBD (in maroon lines). (C) The PRS sensor profiles for the XBB.1 RBD (in orange lines) and XBB.1.5 RBD (in maroon lines). (D) The PRS sensor profiles for the BQ.1 RBD (in orange lines) and BQ.1.1 RBD (in maroon lines). The positions of Omicron mutational sites are indicated by brown-colored filled circles. The convergent mutation sites are indicated by yellow-colored filled diamonds.

The PRS analysis revealed consistent effector peaks corresponding to sites with a high allosteric potential (residues 338-353, 400-406, 420-422, 432-436, 450-456, 501-512) that are conserved across all RBD-ACE2 complexes (Figure 10A,B) The highest peaks corresponded to the hydrophobic RBD core (residues 399-402) and RBD segment (residues 505-512) that connects N501Y and Y505H interfacial positions with the central RBD core (Figure 10A,B). The major allosteric effector clusters were also observed in the functionally important segment (residues 450-456) harboring convergent mutational sites N450 and L452R and connecting the RBD core with the ACE2 binding interface (Figure 10A,B). We also observed that the effector distribution peaks featured higher values for XBB.1.5 (Figure 10A) suggesting that structural stability of this variant may amplify the effector roles of the allosteric hotspots. Of particular interest is a comparison of the effector profiles for XBB.1.5 (Figure 10A) and BQ.1.1 variants (Figure 10B). The overall shape of the effector profiles and the distribution of major peaks are quite similar, indicating that the allosteric networks in both variants are controlled through the same key RBD positions. By projecting positions of convergent mutational sites on the distributions, it can be seen that N450, L452R, R493Q, S494 and especially Y501 residues correspond to major allosteric centers that regulate long-range couplings (Figure 10A,B). There is a visible concentration of the effector sites near the critical interfacial region anchored by Y501 position which is experimentally shown to be involved in short and long-range epistatic relationships with other RBD sites.^64,65,132^

These sites could mediate an ensemble of well-defined signaling paths from the RBD core to the interface regions. Interestingly, by inspecting the effector potential for the entire spectrum of Omicron mutational sites in XBB.1.5 and BQ.1.1 variants we noticed that many of other Omicron positions featured only moderate effector values. The sensor profiles were similar between XBB.1/XBB.1.5 and BQ.1/BQ.1.1 variants (Figure 10C,D) featuring convergent mutation sites N460K and F486S/V as important local peaks of the distribution. It is worth mentioning that highly flexible peripheral RBD regions displayed higher sensor peaks and could be involved in transmission of allosteric changes We argue that these sensor clusters may allow the S-RBD to exploit conformational plasticity to mediate binding response to various classes of antibodies.

We also performed structure-based PRS analysis of major macrostates (Figure 11). Interestingly, we observed significant differences in both the effector and sensor profiles. Consistent with the major contribution of macrostate 4 to the population of XBB.1/XBB.1.5 and BQ.1.1 states, the effector profile for this macrostate featured highest peaks (Figure 11A). Notably, a significant fraction of convergent mutation sites including N440K, K444T, V445P, G446S, N450, and L452R exhibited high effector potential values in macrostate 4. Moreover, the effector potential of these sites diminished in more dynamic macrostates 3 and 6 that contribute significantly to the population distribution of BQ.1 and BQ.1.1 variants (Figure 11A). Another cluster of notable effector centers is concentrated near R493Q, S494 and Y501 sites. On the other hand, highly flexible convergent mutation sites T478R/K, E484K and F486S/P are prominently featured among sensor hotspots of allosteric perturbations (Figure 11B). The sensor potential of these residues is amplified in macrostate 6 in which flexibility of the loops 440-452 and 470-491 is redistributed and enhanced as compared to macrostate 4. Hence, convergent mutations in the RBM regions may function as sensor hotspots of allosteric signals that is enabled through modulation of conformational plasticity in these regions.

**Figure 11.**
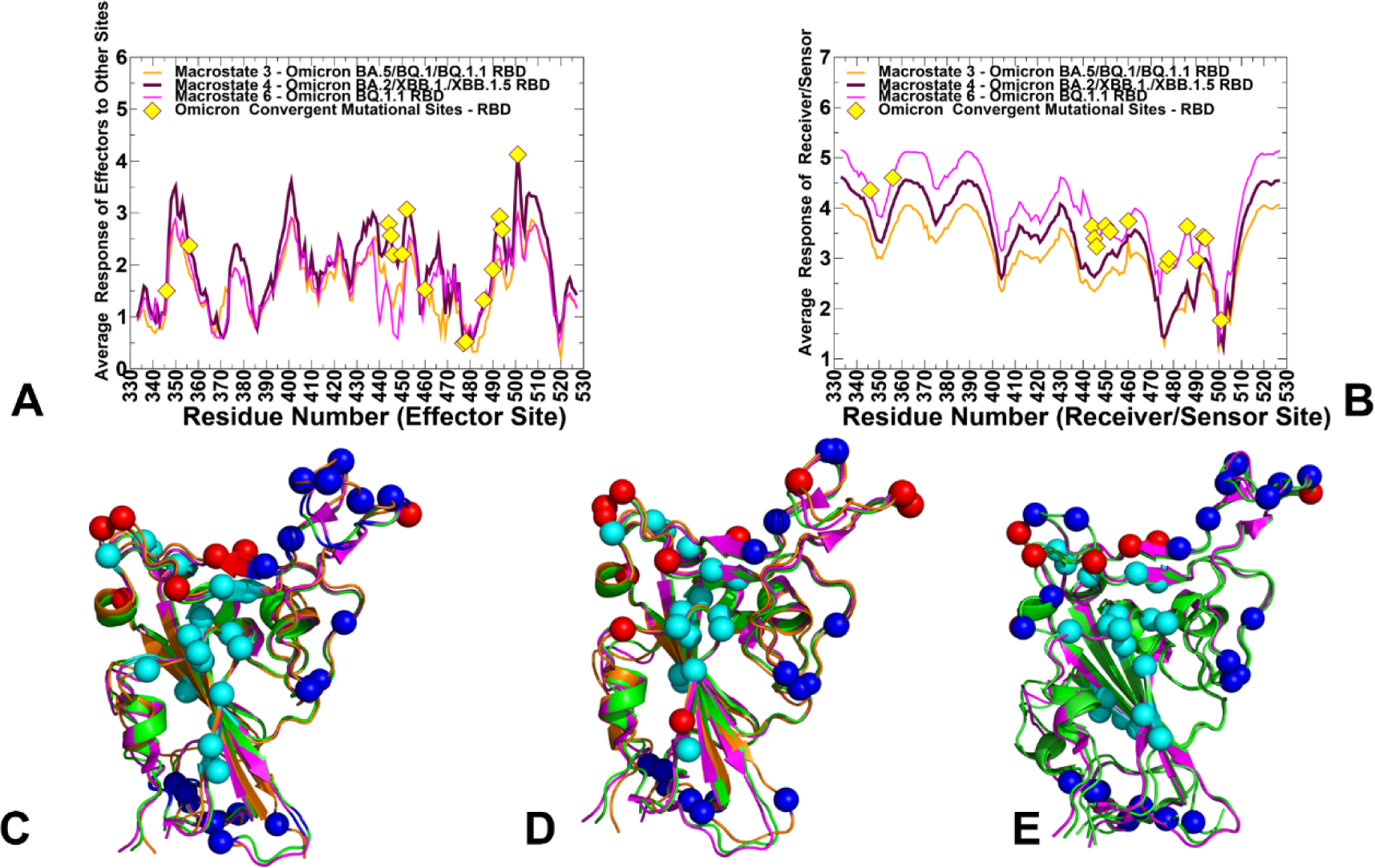
The PRS analysis of the Macrostates for the Omicron RBD-ACE2 complexes. (A) The PRS effector profiles for macrostate 3 (in orange lines), macrostate 4 (in maroon lines) and macrostate 6 (in magenta lines). (B) The PRS sensor profiles for macrostate 3 (in orange lines), macrostate 4 (in maroon lines) and macrostate 6 (in magenta lines).The convergent mutation sites are indicated by yellow-colored filled diamonds. Structural mapping of the allosteric effector and sensor centers in the macrostate 3 (C), macrostate 4 (D) and macrostate 6 (E). The RBD conformations for macrostates are shown in orange ribbons and overlayed with crystallographic BA.2 RBD conformation, pdb id 7XB0 (in green ribbons) and BA.5 RBD conformation, pdb id 7XWA (in magenta ribbons). The allosteric effector centers are shown in cyan spheres and allosteric sensor centers are in blue spheres. The convergent mutation sites are shown in red spheres. Convergent mutation sites that correspond to the effector hotspots (K444T, N450, L452R, Y501) are shown in cyan spheres and convergent mutation sites that correspond to sensor centers (T478R/K, E484K and F486S/P) are accordingly in blue spheres.

Structural superposition of the major macrostates with the crystallographic conformations of BA.2 and BA.5 RBD showed overall fold similarity, while also highlighting differences in modulation of the flexible RBD loops (Figure 11C-E). Macrostate 3 displayed a distorted RBM conformation that abrogated “hook-like” folded arrangement of the RBM tip (Supporting Information Figure S5A). At the same time, macrostate 4 is similar to the crystallographic BA.2 and BA.5 conformations showing a close structural overlap in the loops 440-452 and 470-491 and (Supporting Information Figure S5B). On the other hand, macrostate 6 featured a significant displacement of the loop 440-452 and moderate fluctuations of the RBM tip region (Supporting Information Figure S5C).

Collectively, these observations suggested that convergent mutation sites may play complementary roles in shaping up the long-range allosteric couplings and mediating changes in conformational plasticity of the RBD. Indeed, K444T, N450, L452R and Y501 correspond to the effector centers of allosteric interactions and could control conformational changes in the flexible loop 440-452. The analysis revealed that convergent mutation sites, N460K, T478K, F486/V/S/P of the mobile RBM tip serve as major transmitters of long-range signals in the RBD-ACE2 complexes. According to the PRS analysis, the allosteric cross-talk between sites of convergent mutations acting as allosteric effectors and receivers may provide a mechanism for exerting such coordinated control among interfacial regions. Structural hotspots in adaptation often refer to specific regions or residues that have undergone significant changes or variations during the process of adaptation and which play a critical role in conferring new functions or improving existing functions in response to evolutionary pressures. The observed variations of conformational flexibility in the two flexible RBD loops 440-452 and 470-491 observed in the macrostates, may be evolutionary beneficial for immune escape without compromising binding and stability. Combined, the results of MSM analysis and PRS modeling suggested that convergent mutations may be involved in long-range epistatic interactions where mutational changes in spatially distant communicating residues may control the extent of conformational plasticity in the flexible regions. According to our findings, convergent mutations in the RBD may enable variant-specific adaptation of conformational dynamics in the interfacial regions, and shape up conformational landscapes to control immune escape while the key spike functionality and ACE2 binding are preserved. A potential implication of these results is cumulative effects of individually near-neutral RBD mutations on the conformational landscape may allow for evolutionary adaptation and modulation of conformational mobility,^135^ thereby providing a mechanism for balancing fitness tradeoffs between ACE2 binding and modulation of efficient immune escape.

### Inferring Evolution of Allosteric Cryptic Pockets from Conformational Ensembles and Macrostates: Hidden Role of Convergent Mutation Sites as Modulators of Spike Exposure to Binding Partners

The reversed allosteric communication approach is based on the premise that allosteric signaling in proteins is bidirectional and can propagate from an allosteric to orthosteric site and vice versa.^136-139^ A more integrated computational and experimental strategy exploited the reversed allosteric communication concepts to combine MD simulations and MSM analysis to monitor shifts in the protein conformational ensembles and detect cryptic allosteric sites.^138,139^ We have recently developed a fast and accurate allosteric site prediction method PASSer^88^ and here this approach was employed for detection and monitoring evolution of allosteric pockets in the RBD using conformational ensembles of the Omicron RBD-ACE2 complexes. To simplify the presentation of the results, we focused on the predicted RBD pockets for the optimized structural models of XBB and BQ variants (Supporting Information Figures S6,S7) as well as on detailed analysis of cryptic pockets specific for each of the determined macrostates (Figure 12). The allosteric pockets predicted in the structural models of the XBB and BQ variants shared a conserved and experimentally determined allosteric site in the RBD core where the essential free fatty acid linoleic acid (LA) binds.^140^ The cryo-EM structure of the SARS-CoV-2 S linoleic acid complex revealed this distal binding site and experimentally validated allosteric effect of LA binding. The conserved allosteric pocket discovered experimentally is lined up by residues F338, V341, F342, F377, F374, F392, and W436. Our predicted pocket in this region is formed by residues F338, F342, Y369, F371, A372, I434, and W436 (Supporting Information Figures S6,S7). Notably, these residues also emerged as allosteric effector hotspots in the PRS analysis of allosteric communications in the Omicron complexes. In addition, the analysis also revealed that allosteric pockets may be formed near the RBM region (residues R454, F456, R457, K458, S469, E471, I473) (Supporting Information Figures S6,S7). Interestingly, we also found that BQ.1 and BQ.1.1 RBD structures may open up allosteric pocket that is formed by functional sites near binding hotspot Y501 and convergent mutational hotspots L452R, S494. This pocket was detected only for these Omicron variants and is formed by RBD residues S494, Y495, R403, L452R, Y453, Y501 and G502 (Supporting Information Figure S7). Hence, we suggested that convergent mutational sites owing to evolutionary adaptability and conformational plasticity may be involved in modulation of opening/closing of cryptic allosteric sites. To examine this conjecture in more detail, we also probed allosteric sites using conformational ensembles and reported the PASSer-based pocket predictions of the MSM-derived macrostates (Figure 12). To further assure consistency of PASSer results in detection of allosteric pockets, we also examined macrostates and probed binding pockets using an alternative approach P2Rank/PrankWeb.^141,142^

**Figure 12.**
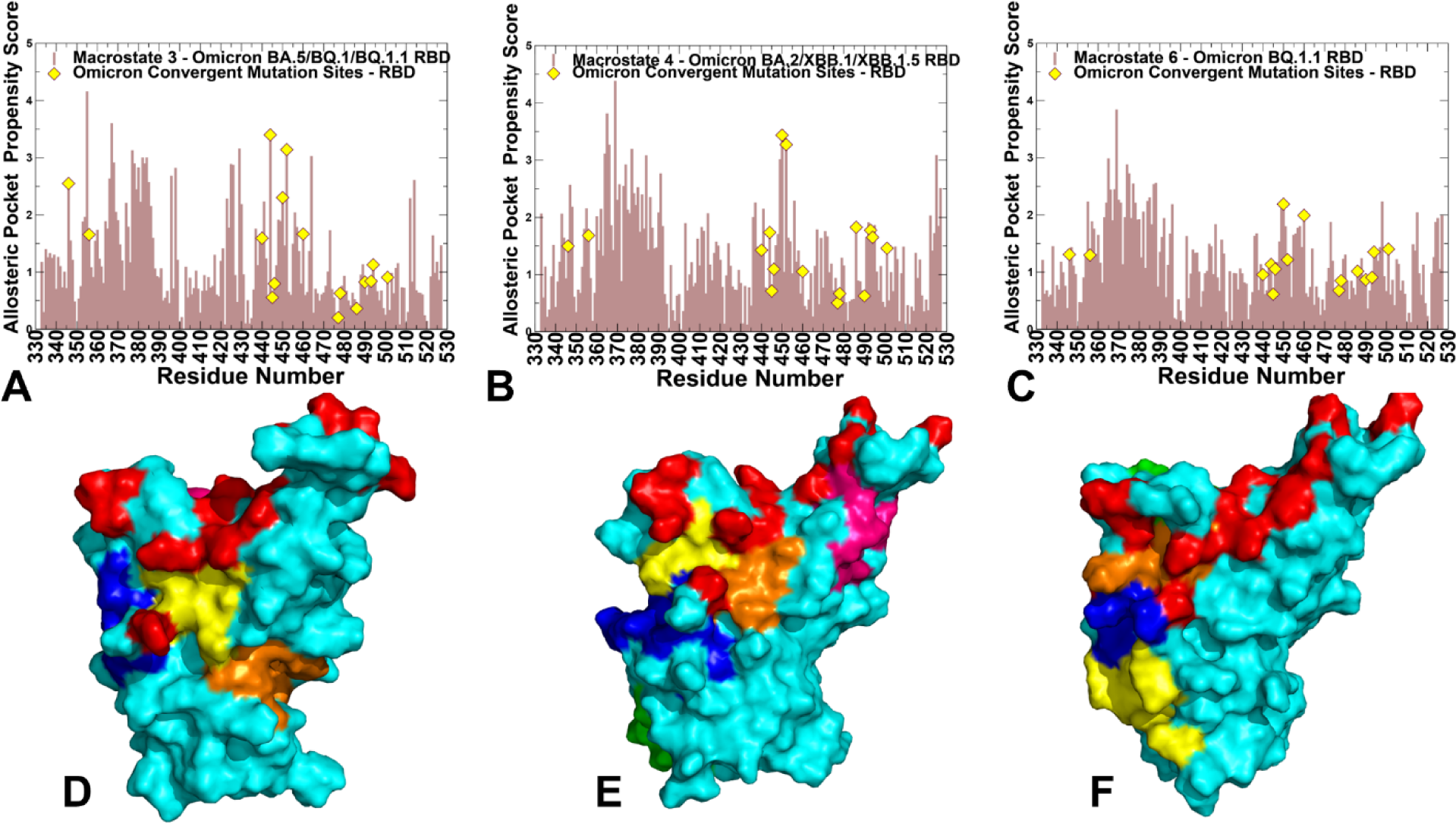
The allosteric pocket detection analysis of the Omicron RBD-ACE2 complexes. The residue-based allosteric pocket propensity scores of the RBD residues computed for representative RBD conformations corresponding to macrostate 3 (A), macrostate 4 (B) and macrostate 6 (C). The allosteric propensity profiles are in light brown bars. The convergent mutation sites (R346, K356, N440, K444, V445, G446, N450, L452, N460, T478, F486, F490, R493 and S494) are depicted in yellow-colored filled diamonds. Structural mapping of PASSer-predicted allosteric pockets on representative RBD conformations corresponding to macrostate 3 (D), macrostate 4 (E) and macrostate 6 (F). The RBD conformations are in cyan surface and convergent mutation sites are shown in red. All macrostates share the predicted RBD pocket (shown in blue surface) that is aligned with the experimentally known allosteric site (residues F338, F342, Y369, F371, A372, I434, and W436). In macrostate 4, the allosteric pocket shown in orange is formed by residues R454, F456, R457, K458, D467, S469, E471, I472 (E). In macrostate 4 the allosteric pocket shown in yellow is formed by residues R346T, T347, N440K, L441, D442, K444, N448, N450 and L452 (E). The RBM pocket for macrostate 4 is shown in pink and consists of residues R454, F456, R457, K458, S469, E471, I473 (E).

We found considerable consistency in recovering the experimentally known allosteric site by using both approaches. By computing the residue-based allosteric pocket propensities^141^ for RBD residues in seven major macrostates we found presence of dense cluster in the RBD core associated with the experimentally known LA-binding site (Figure 12A-C). Hence, this conserved pocket is also preserved in the conformational ensembles and is shared between all Omicron variants. In addition, we also discovered that the distribution peaks can be associated with the RBD regions near the binding hotspot Y501/H505 and strong contributions of convergent mutational sites K444, N450 and L452 (Figure 12A,B). These peaks are particularly strong in macrostates 3 and 4 that are highly prevalent in XBB.1/XBB.1.5 variants. While the composition of these pockets may change in the macrostates it appeared that these functional convergent mutations tend to orchestrate the formation and evolution of cryptic pockets in the conformational ensembles.

Structural mapping showed evolution of the cryptic pockets in the RBD while also highlighting the preservation of the experimentally determined binding site (Figure 12D-F). The results show that variant-specific redistribution of macrostates preserves the experimentally known allosteric pocket on the RBD while allowing for the emergence of hidden allosteric pockets that are anchored by convergent mutation sites K444T and L452R. In the macrostate 3 with a disordered RBM tip, the allosteric pocket near the RBM region may “dissolve” providing a very plastic and “pocketless” interface in this region (Figure 12D). Similarly, macrostate 6 featuring altered and highly flexible loop 440-452 may induce stabilization of the RBD core pockets that occupy regions near the experimental allosteric site (Figure 12F). Some of these pockets have a transient character and may emerge and disappear in different conformational states. In the dominant macrostate 4, we found several conserved pockets including the experimentally discovered allosteric site along with two adjacent small pockets and RBM pocket (Figure 12E). These allosteric binding sites are maintained in the dominant macrostate and could be targeted in both XBB and BQQ variants. One of the more interesting observations of the allosteric pocket analysis was detection of two additional pockets in the macrostate 4 (shown in orange and yellow on Figure 12E) that are adjacent to the experimental allosteric site and contain a number of convergent mutation sites. In particular, one of these pockets (shown in yellow on Figure 12E) is formed by residues R346T, T347, N440K, L441, D442, K444, N448, N450 and L452. Strikingly, this allosteric pocket included several key convergent mutation sites (R346T, K444, N450 and L452) that also play role as allosteric effector centers. Another pocket shown in orange on Figure 12E (residues R454, F456, R457, K458, D467, S469, E471, I472) links the RBD core with the binding interface. Both these allosteric pockets are anchored by convergent mutation sites N450 and L452R. Hence, these hotspots could play an integrated role in modulating conformational plasticity and viability of these pockets. Together, these results showed that a selective group of convergent mutation sites could function as allosteric effector centers that control evolution and distribution of allosteric pockets through modulation of conformational plasticity in the flexible adaptable regions of the RBD. These findings indicated that Omicron XBB.1.5 and BQ.1.1 subvariants may evolve to accumulate convergent mutations in flexible sites to modulate allostery and immune escape through opening/closing patches/pockets to control access of antibodies.

## Conclusions

The new generation of SARS-CoV-2 Omicron variants displayed a significant growth advantage and the increased viral fitness by acquiring convergent mutations in the N-terminal and receptor binding domains of spike proteins, suggesting that the immune pressure can promote convergent evolution leading to the observed sudden acceleration of SARS-CoV-2 evolution. Evolutionary and functional studies explored various scenarios underlying balance of virus fitness tradeoffs, but a systematic analysis of structural and dynamics effects of convergent mutations in newly emerged wave of Omicron variants is conspicuously lacking. In the current study, we combine extensive microsecond MD simulations and Markov state models to characterize conformational landscapes and identify specific dynamic signatures of the SARS-CoV-2 spike complexes with the host receptor ACE2 for the recently emerged highly transmissible XBB.1, XBB.1.5, BQ.1, and BQ.1.1 Omicron variants. Microsecond simulations and Markov modeling provided a detailed characterization of the conformational landscapes and revealed the increased thermodynamic stabilization of the XBB.1.5 subvariant which is contrasted to more dynamic BQ.1 and BQ.1.1 subvariants. Despite considerable structural similarities, Omicron mutations can induce unique dynamic signatures and specific distributions of conformational states that explain the patterns of stability and immune escape in the Omicron variants. Using a comparative MSM analysis we showed that the distribution of dominant macrostates in the Omicron variants can allow for allosteric modulation of mobility in the flexible RBD loops 470-491 and 440-452 involved in the interfacial contacts and containing critical convergent mutation sites. Our results suggested that variant-specific changes of conformational mobility in these functional loops can be fine-tuned through cross-talk between convergent mutations thereby providing an evolutionary path for the improved immune escape without significant cost on the ACE2 binding. By combining MD simulations and MSM analysis with perturbation-based response scanning approaches, we also examined mechanisms of long-range dynamic couplings and allosteric communications in the Omicron RBD-ACE2 complexes. Based on premise that allostery is linked with the evolvability of proteins, we explored the relationship between allosteric interactions and patterns of convergent Omicron mutations, showing how convergent mutations can potentiate conformational plasticity and modulate allosteric responses in the RBD to binding and immune escape. Through perturbation response scanning analysis, our study reveals important complementary roles of convergent mutation sites as effectors/regulators and sensors/receivers of allosteric signal transmission in the RBD-ACE2 complexes. We also explored the reversed allosteric communication approach and characterized the effect of dynamics on the distribution of allosteric pockets in the Omicron subvariants. The results show that variant-specific redistribution of macrostates preserves the experimentally known allosteric pocket on the RBD while allowing for the emergence of hidden allosteric pockets that are anchored by convergent mutation sites K444T and L452R. The findings support a mechanism according to which Omicron mutations may have evolved to balance thermodynamic stability and conformational adaptability in order to ensure proper tradeoff between stability, binding and immune escape.

## Author Contributions

Conceptualization, G.V. and P.T.; methodology, S.X., G.V. P.T.; software, S.X., M.A., G.G., G.V and P.T.; validation, G.V.; formal analysis, G.V., M.A., G.G., S.X., and P.T. investigation, G.V. and P.T.; resources, G.V., M.A. and G.G.; data curation, G.V.; writing—original draft preparation, P.T. and G.V.; writing—review and editing, S.X., G.V., M.A. and G.G.; visualization, G.V.; supervision, G.V.; project administration, P.T. and G.V.; funding acquisition, P.T. and G.V. All authors have read and agreed to the published version of the manuscript.

## Data Availability Statement

Data is fully contained within the article and Supplementary Materials. Crystal structures were obtained and downloaded from the Protein Data Bank (http://www.rcsb.org). All simulations were performed using openMM high-performance toolkit for molecular simulation that was obtained from websites https://openmm.org/; https://simtk.org/projects/openmm and https://github.com/openmm/openmm. All simulations were performed using the all-atom additive CHARMM36 protein force field that can be obtained from http://mackerell.umaryland.edu/charmm_ff.shtml. The rendering of protein structures was done with UCSF ChimeraX package (https://www.rbvi.ucsf.edu/chimerax/) and Pymol (https://pymol.org/2/). The software tools used in this study are freely available GitHub sites https://github.com/smu-tao-group/protein-VAE; https://github.com/smu-tao-group/PASSer2.0. All the data obtained in this work (including simulation trajectories, topology and parameter files, the software tools, and the in-house scripts are freely available at DOI 10.5281/zenodo.7942171. Version 1 10.5281/zenodo.7942172.

## Acknowledgments

G.V acknowledges support from Schmid College of Science and Technology at Chapman University for providing computing resources at the Keck Center for Science and Engineering.

