## Supplemental Figures S1,S2, S3, S4, S5, S6, S7 for "Markov State Models and Perturbation-Based Approaches Reveal Distinct Dynamic Signatures and Hidden Allosteric Pockets in the Emerging SARS-Cov-2 Spike Omicron Variants Complexes with the Host Receptor: The Interplay of Dynamics and Convergent Evolution Modulates Allostery and Functional Mechanisms"

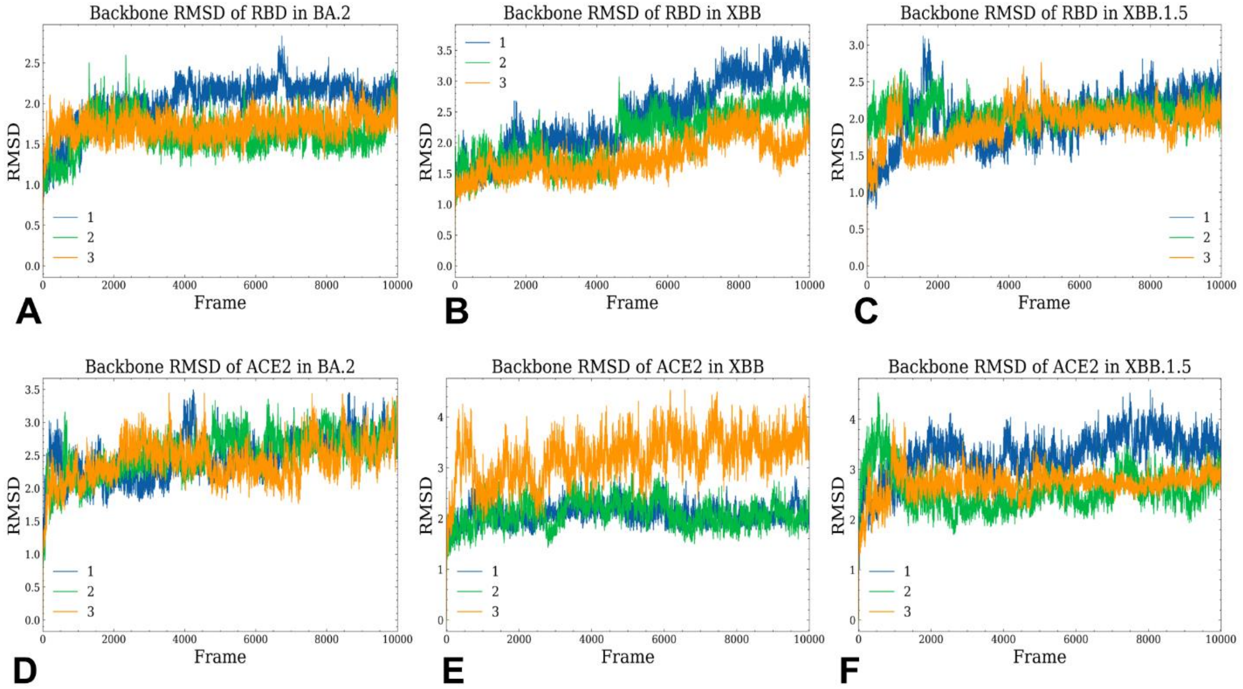


**Figure S1.** Conformational dynamics profiles obtained from 3 microsecond-long all-atom MD simulations of the Omicron RBD BA.2, XBB.1 and XBB.1.5 complexes with hACE2. The RMSD profiles for the RBD residues obtained from 3 microsecond MD simulations of the Omicron RBD BA.2-ACE2 complex, pdb id 7XB0 (A), Omicron RBD XBB.1-ACE2 complex (B), and Omicron RBD XBB.1.5-ACE2 complex (C). The RMSD profiles for ACE2 residues for the Omicron RBD BA.2-ACE2 complex (D), Omicron RBD XBB.1-ACE2 complex (E), and Omicron RBD XBB.1.5-ACE2 complex (F).

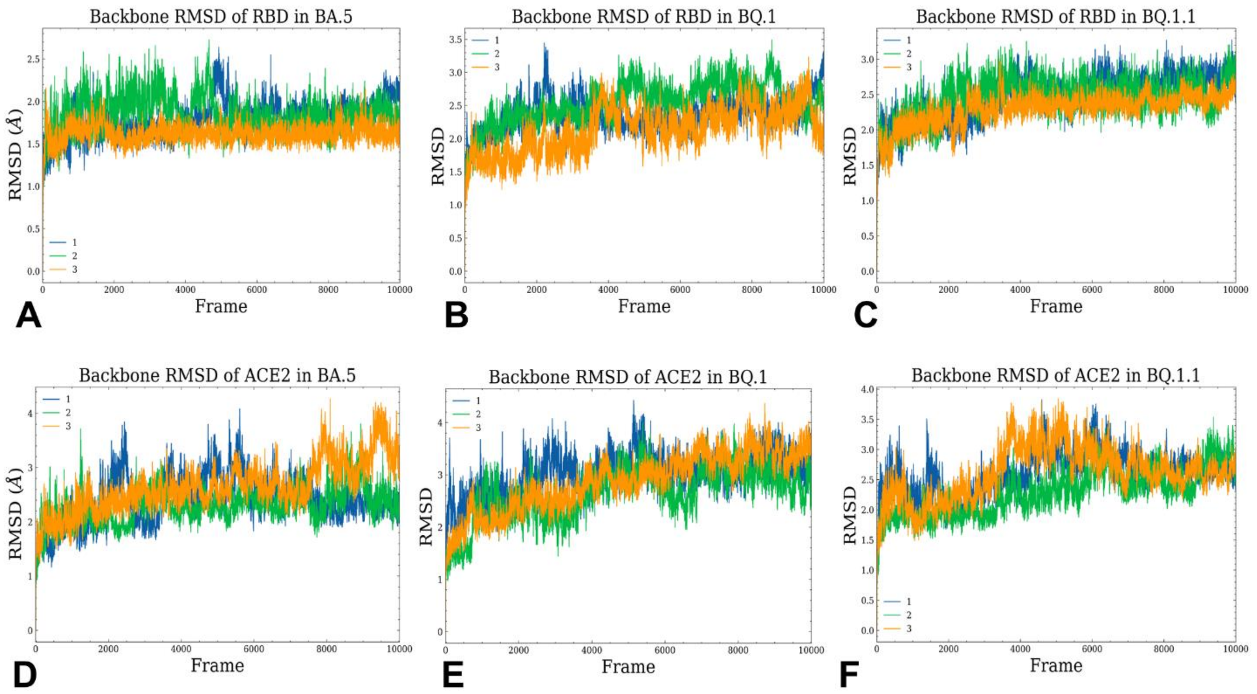


**Figure S2.** Conformational dynamics profiles obtained from 3 microsecond-long all-atom MD simulations of the Omicron RBD BA.5, BQ.1 and BQ.1.1 complexes with hACE2. The RMSD profiles for the RBD residues obtained from 3 microsecond MD simulations of the Omicron RBD BA.5-ACE2 complex, pdb id 7XWA (A), Omicron RBD BQ.1-ACE2 complex (B), and Omicron RBD BQ.1.1-ACE2 complex (C). The RMSD profiles for ACE2 residues for the Omicron RBD BA.5-ACE2 complex (D), Omicron RBD BQ.1-ACE2 complex (E), and Omicron RBD BQ.1.1-ACE2 complex (F).

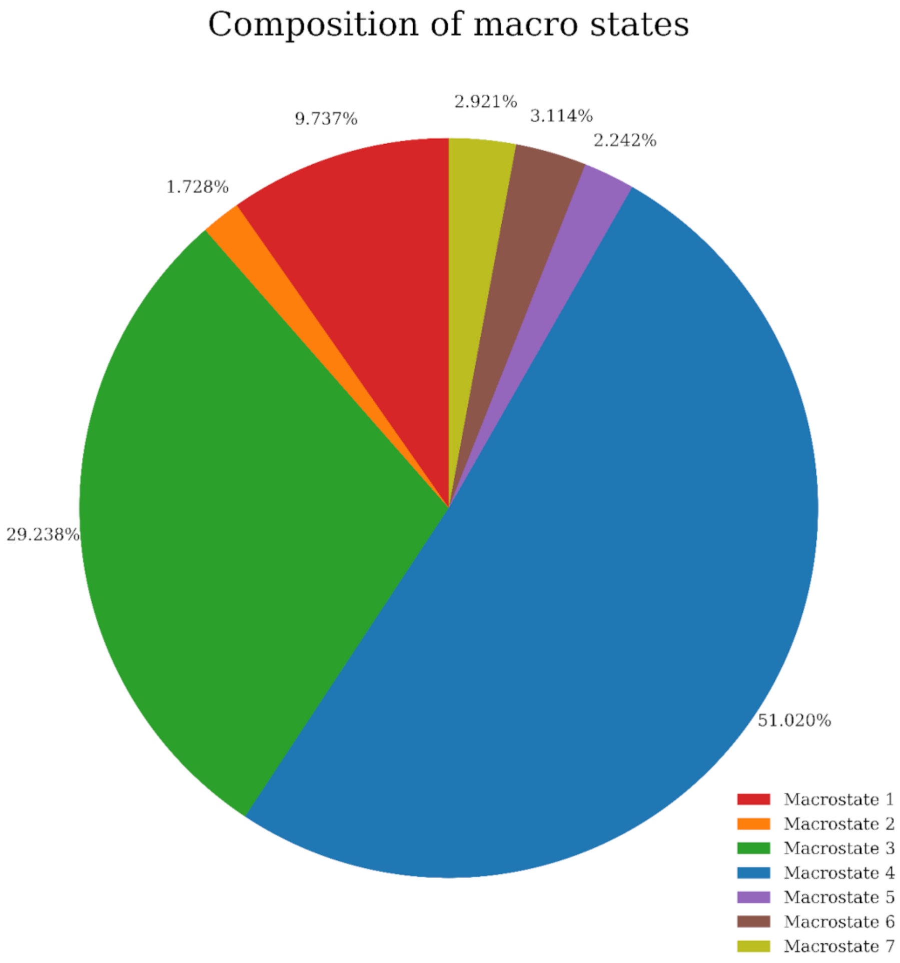


**Figure S3.** The pie diagram of the cumulative population of different macrostates in the studied Omicron RBD-ACE2 complexes.

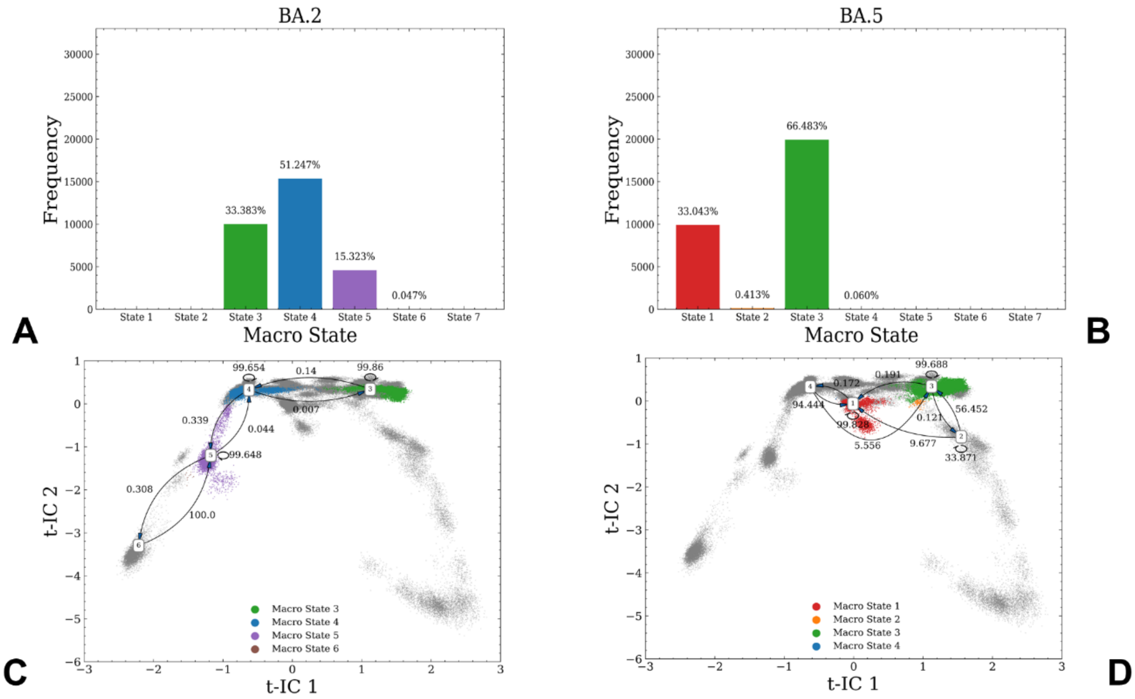

**Figure S4.** MSM analysis of the conformational landscape for the parental BA.2 and BA.5 RBD-ACE2 complexes. The frequency distribution of occupying different macrostates in BA.2 (A) and BA.5 (B). The transition probability maps among different macrostates with 3 ns lag time for BA.2 (C) and BA.5 variant (D). The high percentage of self-conserved probability shows the stability of macrostates.

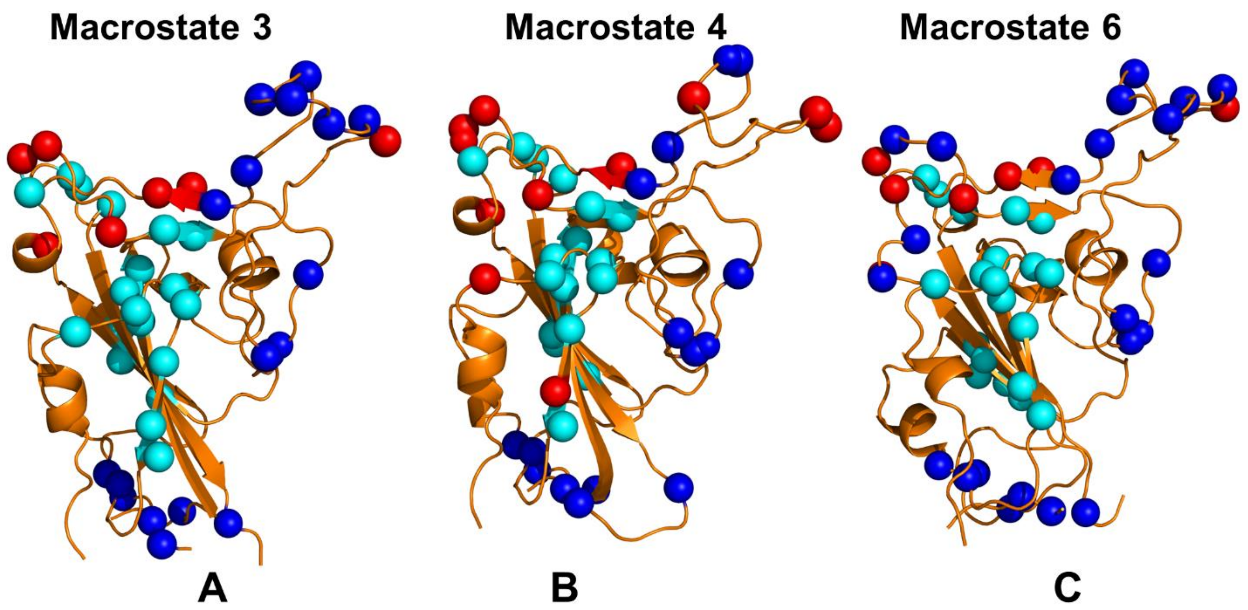


**Figure S5.** Structural mapping of the allosteric effector and sensor centers in the macrostate 3 (C), macrostate 4 (D) and macrostate 6 (E). The RBD conformations for macrostates are shown in orange ribbons. The allosteric effector centers are shown in cyan spheres and allosteric sensor centers are in blue spheres. The convergent mutation sites are shown in red spheres. Convergent mutation sites that also correspond to the effector hotspots (K444T, N450, L452R, Y501) are shown in cyan spheres and convergent mutation sites that correspond to sensor centers (T478R/K, E484K and F486S/P) are accordingly in blue spheres.

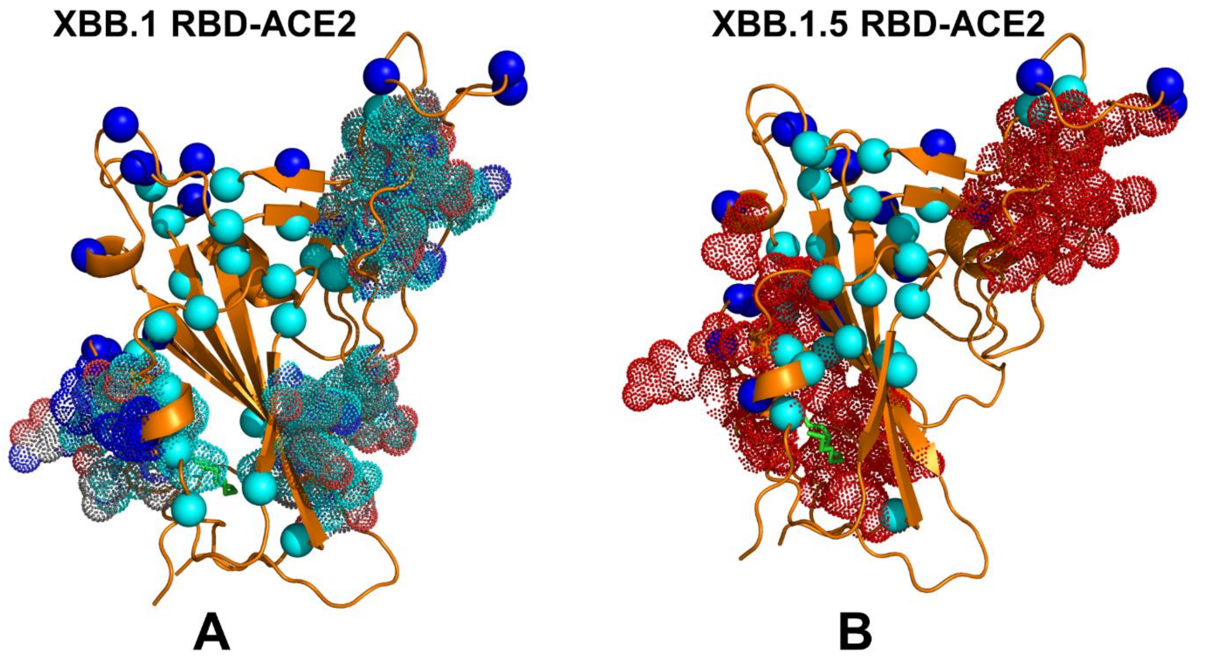


**Figure S6.** Structural mapping of the allosteric hotspots and allosteric pockets in the Omicron XBB.1 (A) and XBB.1.5 RBD complexes. The RBD conformations are shown in orange ribbons. The allosteric effector hotspots are shown in cyan spheres. Omicron mutation sites are shown in blue spheres. The PASSer-predicted allosteric pockets are highlighted in dot spheres.

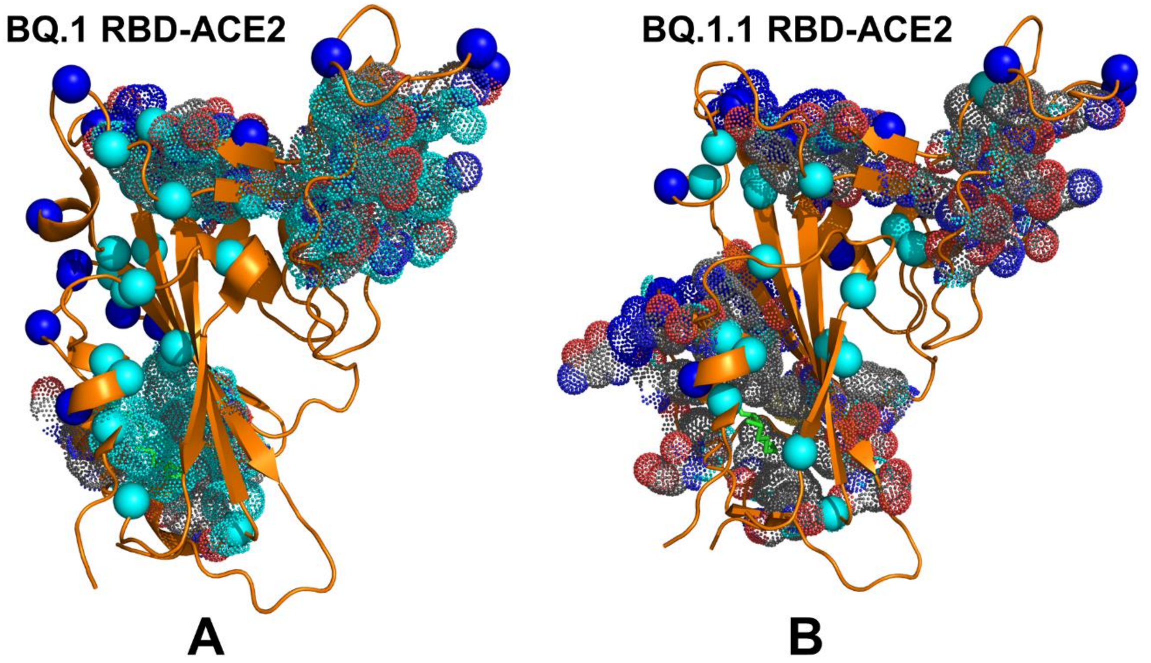


**Figure S7.** Structural mapping of the allosteric hotspots and allosteric pockets in the Omicron BQ.1 (A) and BQ.1.1 RBD complexes. The RBD conformations are shown in orange ribbons. The allosteric effector hotspots are shown in cyan spheres. Omicron mutation sites are shown in blue spheres. The PASSer-predicted allosteric pockets are highlighted in dot spheres.
